# Designing patient-oriented combination therapies for acute myeloid leukemia based on efficacy/toxicity integration and bipartite network modeling

**DOI:** 10.1101/2023.05.25.542238

**Authors:** Mehdi Mirzaie, Elham Gholizadeh, Juho J. Miettinen, Filipp Ianevski, Tanja Ruokoranta, Jani Saarela, Mikko Manninen, Susanna Miettinen, Caroline A. Heckman, Mohieddin Jafari

## Abstract

Acute myeloid leukemia (AML), a heterogeneous and aggressive blood cancer, does not respond well to single-drug therapy. A combination of drugs is required to effectively treat this disease. Computational models are critical for combination therapy discovery due to the tens of thousands of two-drug combinations, even with approved drugs. While predicting synergistic drugs is the focus of current methods, few consider drug efficacy and potential toxicity, which are crucial for treatment success. To find effective new drug candidates, we constructed a bipartite network using patient-derived tumor samples and drugs. The network is based on drug-response screening and summarizes all treatment response heterogeneity as drug response weights. This bipartite network is then projected onto the drug part, resulting in the drug similarity network. Distinct drug clusters were identified using community detection methods, each targeting different biological processes and pathways as revealed by enrichment and pathway analysis of the drugs’ protein targets. Four drugs with the highest efficacy and lowest toxicity from each cluster were selected and tested for drug sensitivity using cell viability assays on various samples. Results show that the combinations of ruxolitinib-ulixertinib and sapanisertib-LY3009120 are the most effective with the least toxicity and best synergistic effects on blasts. These findings lay the foundation for personalized and successful AML therapies, ultimately leading to the development of drug combinations that can be used alongside standard first-line AML treatment.

**Key Points:**

- Ruxolitinib-ulixertinib and sapanisertib-LY3009120 have the best synergistic effects on AML, with the least toxicity.
- This study’s combinations destroy blasts without harming other healthy cells, unlike standard chemotherapy, which is less specific.

## Introduction

Acute myeloid leukemia (AML) is an inter-and intra-tumor heterogeneous disease ^1,2^. It is identified when the bone marrow (BM) contains at least 20% of blast cells of the myeloid lineage ^3^. Traditional chemotherapeutics have limited efficacy in patients over the age of 65, with a survival rate of less than 25% at one year follow-up and less than 9% after 5 years ^4^. Despite recent advances in genome sequencing, which enables researchers to identify a large number of mutations, we are still hampered by the absence of drugs which specifically target cancer-associated mutations ^5^. On the other hand, the majority of AML patients do not have actionable mutations, and the link between cancer genotype, phenotype, and therapeutic function is poorly understood ^6^. Even if we overcome the above difficulties and identify the exact mutations in genotype, monotherapy drug resistance will remain a major clinical complication ^7^. Targeted anti-cancer compounds used in combination therapy have the potential to overcome resistance, improve patient response to current treatments, reduce dose-limiting single agent toxicity, and broaden the spectrum of available therapies ^8^.

Drug combination therapy offers the chance to suppress a number of pathways synergistically, including patient-specific cancer rescue pathways and phenotypic redundancy across heterogeneous cancer sub-clones ^9^. The phenotypic effects of thousands of drug combinations can be evaluated in patient-derived cells and other pre-clinical model systems using high-throughput screening. However, because there are so many possible drug and dose combinations, large-scale multi-dose combinatorial screening is not recommended, due to the limited number of cells available from patient samples. Using the presented method in this study, researchers would be able to categorize the most important AML drugs into different clusters, each of which target proteins associated with various signaling pathways.

In our earlier research, we designed a systems pharmacology approach based on network modeling to identify prospective drug combinations in AML ^10^. To gain a deeper understanding of the factors that govern drug response in AML patients, we utilized a unique and extensive dataset obtained through drug response screening of samples from both AML patients and healthy donors in Finland as part of our study. Our model accounts for the efficacy and toxicity of drug response, which are simultaneously evaluated on patient and healthy samples, respectively ^11^. A weighted bipartite network composed of two parts, chemical components, and patient samples was built to develop a drug combination strategy using the screening outcomes of single drug responses on AML patient samples. This enables researchers to directly access the phenotype of the patients’ cancer cells through *ex vivo* drug response data, and by using network modeling and clustering analysis, demonstrate the drugs functionalities. Next, top drug combinations can be predicted based on phenotypic responses of samples in each cluster. In addition, we used two different computational resources, i.e., molecular biology annotations, and the chemical structure of drugs, to perform intra-cluster homogeneity analysis. The top combinations for further study in current pre-clinical drug combination screening are frequently chosen based on synergy alone, without taking into account the combination efficacy and toxicity effects, despite the fact that these are critical determinants of a therapy’s clinical success ^12^.

Considering the importance of toxicity, in this study, we investigated the drug response of AML patient and healthy donor samples to calculate both efficacy and toxicity, respectively. The efficacy and toxicity of the drugs was assessed *ex vivo* on AML patient samples using cell viability and flow cytometry assays. Since clinical symptoms in patients are caused by blast cell accumulation in bone marrow ^13^, based on the assessment of CD marker expressions, we suggested the most blast-specific combinations as promising combinations for AML treatment. The novel combinations indicated in this study are effective on blasts but have a reduced level of toxicity on other cell populations, whereas with standard chemotherapy the majority of the blasts are killed together with other cell types.

## Results

The entire workflow of this study is depicted in Figure 1 and detailed material and method is presented in the supplementary information. The drug responses of 525 chemical compound tested on 199 AML patients’ bone marrow samples were obtained from the FIMM AML data set^11^. The bipartite network was constructed using this data set, as explained in the materials and methods section in the supplement. A bipartite network can be projected into two different types of unipartite networks containing nodes of only one type. The projection of the bipartite network, onto the “drug” node set is considered here, called the drug similarity network. The Louvain community detection approach was used to find drugs that behaved similarly in terms of drug response^14^. The results gave us two communities (clusters) of drugs denoted by *C*_1_ and *C*_2_ with network sizes of 155 and 141, respectively (Table S1).

**Fig. 1.**
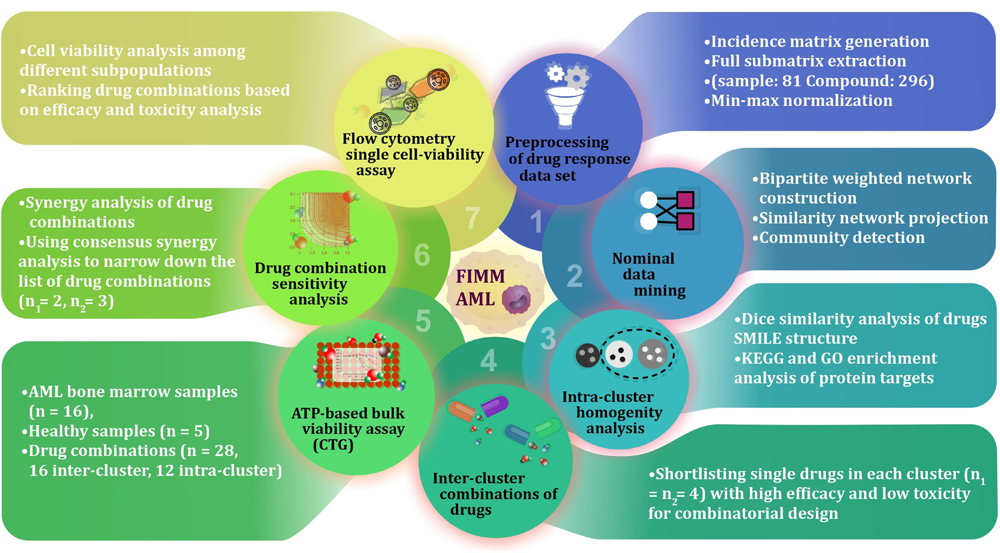
Schematic outline of the study. Data pre-processing began after data collection, which was followed by full matrix extraction, weighted bipartite network reconstruction, and computational validation. After the selection of the best combinations, bone marrow and peripheral blood samples from both healthy individuals (n=5) and AML patients (n= 16) were subjected to drug sensitivity assessment. For ATP-based viability assay the study design contains 8 drugs and 28 combinations in 384-well plates, each drug with 5 different concentrations and two replicate. The single cell sensitivity assay using the iQue® Screener PLUS flow cytometer was performed in 384-well plates to monitor drug effects on cell sub-types. The study design contains 5 drugs and 3 combinations, all with two replicates and five concentrations. For sapanisertib, the drug concentrations are 0.1, 1, 10, 100, and 1000 nM, and for all other drugs are 1, 10, 100, 1000, and 10000 nM.

### Comparing AML drug clusters: Evaluating protein target pathways and chemical structure similarity

We used two independent computational methods to determine how distinct the two clusters are: the first identifies the significant difference between biological pathways of drug protein targets in each cluster, and the second evaluates the chemical structure similarity of drugs in each cluster. We constructed a drug-target network using the drug target commons (DTC) database^15^, which is also a bipartite network in which each link connects drugs to their protein targets. Let *T*_1_ and *T*_2_ represent the set of protein targets of drugs in the cluster *C*_1_ and *C*_2_, respectively, and *T* represents the union of *T*_1_and *T*_2_. In this study |^T^l| = 921, |^T^2| = 842, and |T| = 1055.

Proteins with a score S (explained in the supplementary information 1.3) greater than log (2) are considered to be preferentially targeted by drugs in cluster 1, denoted by PPT1. Similarly, PPT2 proteins have a score of less than *log*(0.50). We performed GSEA (gene set enrichment analysis) on PPT1 and PPT2 proteins based on their associated scoring functions. As expected, the biological processes and signaling pathways affected by drugs in Clusters 1 and 2 are distinct. This difference enables us to inhibit two different signaling pathways using one combination. Drugs in cluster 1 (PPT1), such as LY3009120 (a Pan-RAF inhibitor), predominantly target proteins associated with the RAF-MEK-ERK signaling pathway. This pathway plays a crucial role in cell proliferation and growth, indirectly influencing processes like cell-substrate adhesion and ion trans-membrane transport, which are enriched in our analysis ^16^. In contrast, JAK1/2 inhibitors like ruxolitinib target JAK proteins, involved in cytokine signaling and immune responses, impacting pathways related to neuroactive ligand-receptor interactions and the regulation of actin cytoskeleton ^17^. Drugs like birabresib, which target proteins in the bromodomain and extra-terminal (BET) family, have a role in gene regulation through chromatin binding, affecting gene expression and pathways related to chemical reactions and collagen metabolism ^18^. Plicamycin, which binds to guanine-cytosine-rich regions of DNA, may influence gene expression and regulation, impacting pathways related to collagen metabolism and other DNA-dependent processes (Figure 2 A) ^19^. On the other hand, proteins targeted by drugs in cluster 2 (PPT2) (silmitasertib, ulixertinib, sapanisertib, and teniposide) are in the p53 signaling pathway, cell cycle, apoptosis, and pancreatic, colorectal and chronic myeloid leukemia cancers and related to tumorigenesis and progression pathways, including human immunodeficiency virus 1 infection ^20–23^.

**Fig. 2.**
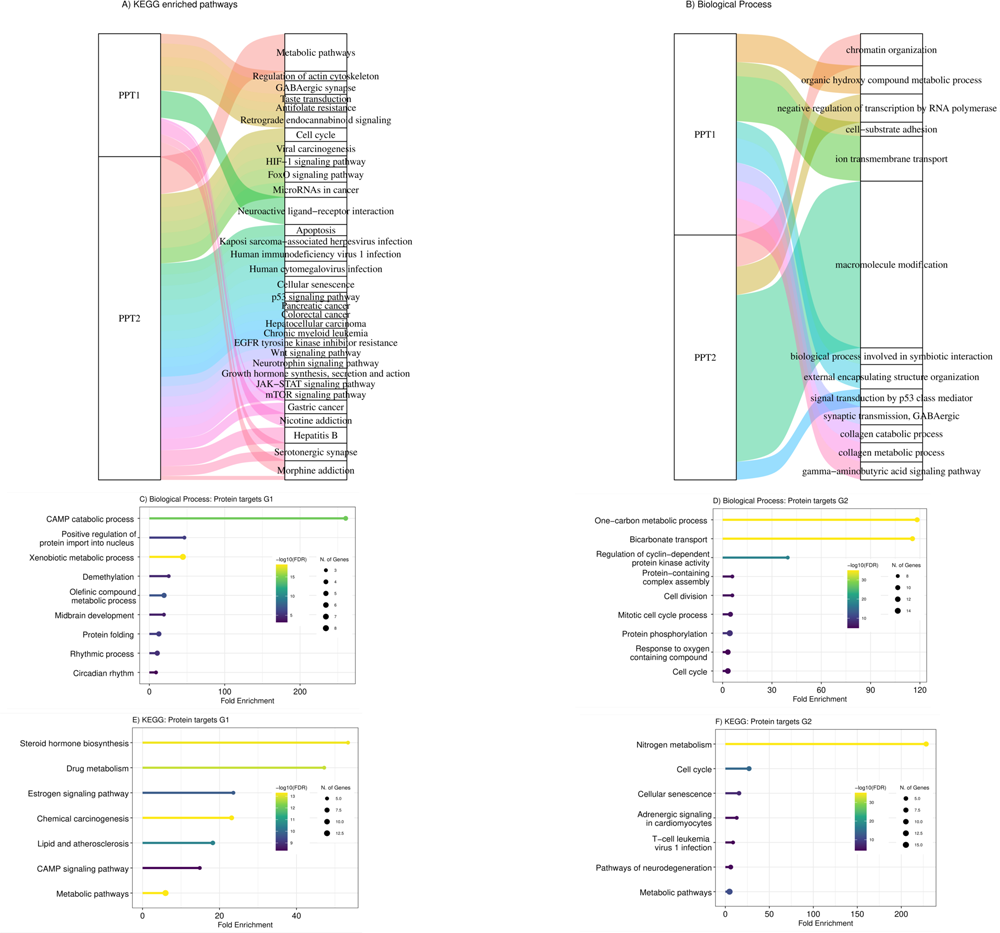
Sankey plot of enriched (A) KEGG signalling pathways and (B) GO biological processes related to target protein clusters PPT1 and PPT2. Each rectangle on the right side represents a pathway or biological process, and the size of each rectangle illustrates the degree of connectivity of each pathway. Each biological process or pathway is represented by a unique color. GO and KEGG pathway enrichment analysis on proteins that are merely targets by drugs in one cluster. G1 (G2) includes proteins that are targeted by at least three drugs in cluster1 (cluster2) (155 and 141 drugs). (C) Biological processes (BPs) of G1, (D) Biological processes (BPs) of G2, (E) KEGG pathway related to G1 proteins and (F) KEGG pathway related to G2 proteins. The size of the node corresponds to number of genes, the x-axis is Fold Enrichment and the color of bars indicates the negative logarithm of Fold Enrichment.

We also performed ShinyGO ^24^ Gene Ontology and KEGG pathway enrichment analysis on proteins that are merely targeted by drugs in one cluster. For this purpose, two protein sets G1 and G2 were selected such that G1 includes proteins targeting by at least three drugs in cluster 1 and at most two drugs in cluster 2, and similarly, G2, consist of proteins that are mostly targeted by drugs in cluster 2 (at least three drugs in cluster 2 and at most two drugs in cluster 1). REVIGO was also used to summarize the enriched GO terms, and the results are shown in Figure 2 and Tables S2 and S3. The cAMP signaling pathway, lipids and atherosclerosis, steroid hormone biosynthesis, and rhythmic processes and circadian rhythm are biological processes related to G1 proteins, which are mostly targeted by LY3009120, birabresib, plicamycin, and ruxolitinib. Cell cycle, cellular senescence, T-cell leukemia virus 1 infection and cell division, mitotic cell cycle, and protein phosphorylation processes are related to G2 proteins, mostly targeted by silmitasertib, ulixertinib, sapanisertib, and teniposide. Therefore, we demonstrate that the protein targets of drugs in each cluster are on different pathways and biological processes. To do homogeneity analysis of chemical structure of drugs, the dice similarity test was used to show how structurally similar the drugs are in each cluster. This measurement compares the number of chemical features shared by a pair of compounds to the average size of the total number of features present. Pairwise similarities were calculated for chemical compounds chosen from two drug clusters for inter-cluster comparison. Drugs from different clusters are less similar than drugs from the same cluster, as shown in the Figure 3A. According to the box plot, the inter-cluster similarities are less than the intra-cluster similarities in both clusters. The results of the t-test imply that the mean of inter-cluster similarities is less than the mean of intra-cluster similarities in clusters 1 and 2 (p-value < 2.2e-16 for both t-test).

**Fig. 3.**
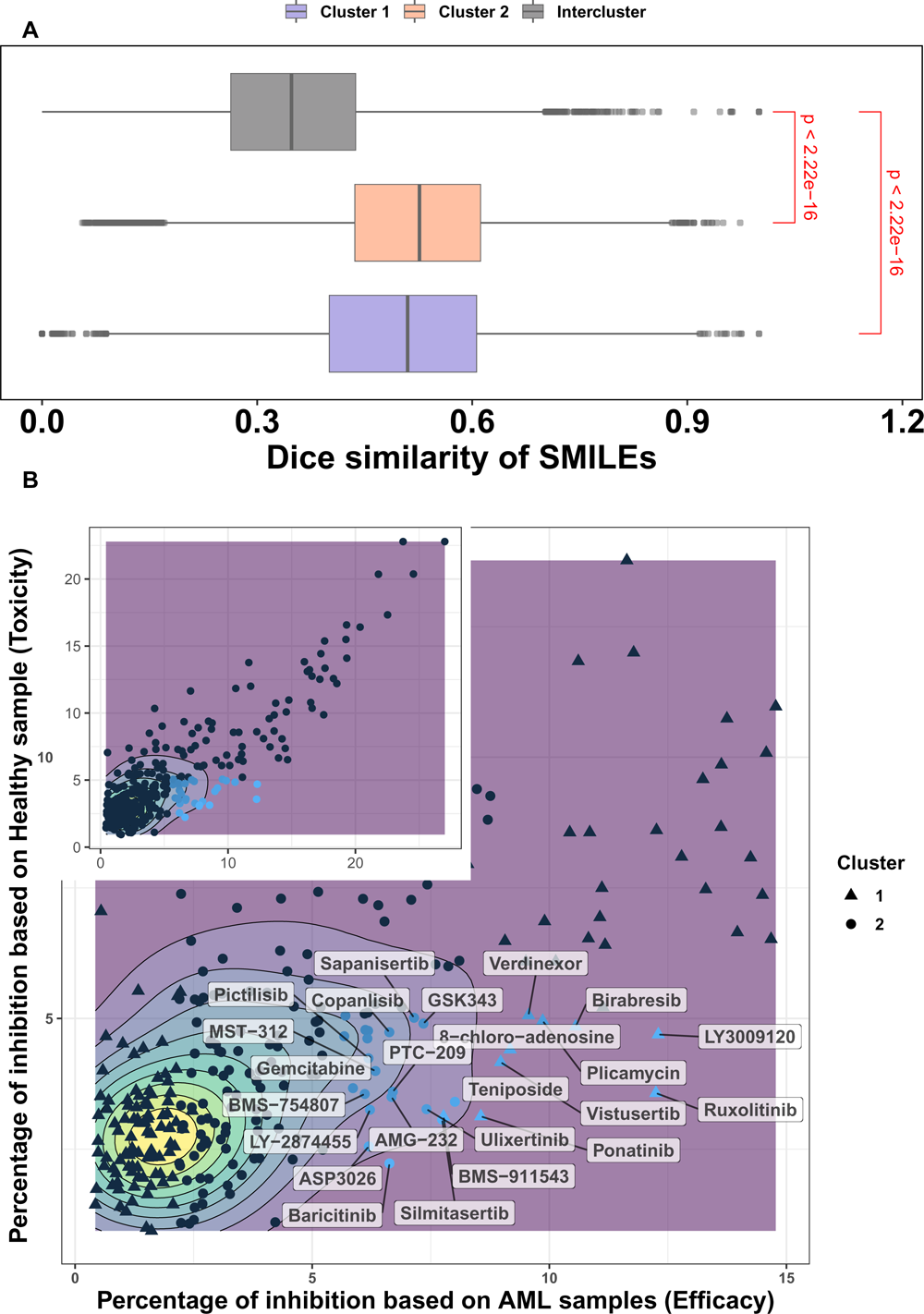
Comparative analysis of dice similarity and drug efficacy-toxicity profiles in AML therapy. (A)Box plot of dice similarity coefficient indices comparing intra-cluster 1 and 2 to inter-cluster compound pairs. P-value is generated using Wilcoxon signed-rank test, shown in red color. (B) The toxicity and efficacy of 296 drugs. Inset plot shows the relationship between toxicity and efficacy. Top five percent of drugs whose toxicity is less than the average of a**l** drug toxicity and whose efficacy is greater than the average of all drug efficacy are in blue, and their name is shown in rectangle labels.

### Combination selection: Balancing toxicity and efficacy across clusters

As a result, we demonstrated that clusters are well-separated and that the protein targets of drugs in each cluster are involved in distinct pathways. In this novel combination strategy, we aim to select two drugs from distinct clusters while taking both toxicity and efficacy into account. The optimal combinations are those that have lower toxicity than the average toxicity and higher efficacy values than the average efficacy value for all drugs. For each drug, the average drug response of healthy and AML patient samples in the data set are considered as toxicity and efficacy, respectively. We assume that the ideal drugs have no inhibitory effect on healthy samples but significantly influence blast cells in AML patient samples. We chose the top 5% of drugs whose toxicity is less than the average of all drug toxicity and efficacy is greater than the average of all drug efficacy. Figure 3B depicts the link between toxicity and efficacy values of 296 drugs on 81 samples. The top selected small molecule in each cluster is summarized in Table 1 and Table S4. Four chemical compounds from cluster 1 including birabresib, LY3009120, plicamycin, and ruxolitinib as well as four drugs from cluster 2 including sapanisertib, silmitasertib, teniposide, and ulixertinib were chosen for drug combination testing. According to our experimental design, the combination of drugs within a single cluster is known as negative group or intra-cluster, and the combination of drugs between clusters is considered as positive group or inter-cluster.

**Table 1.**
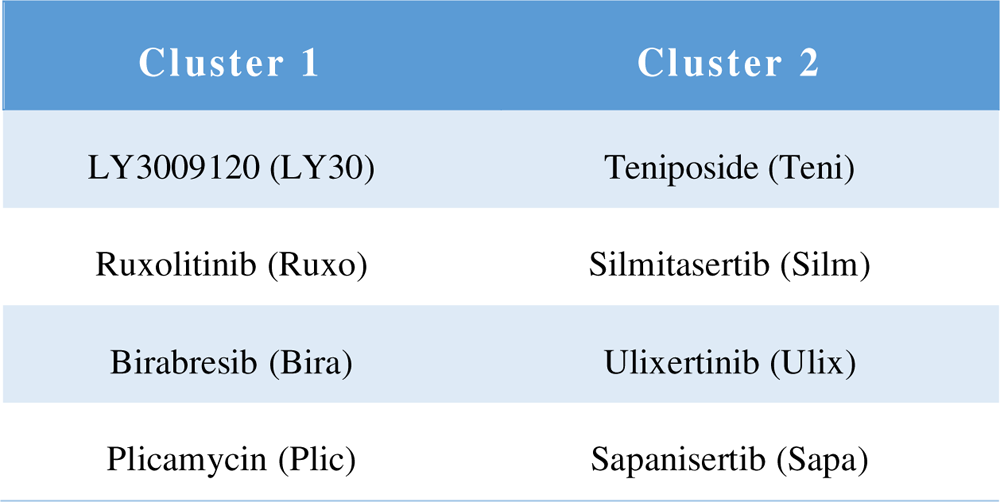
The selected chemical compounds from two clusters of drugs in the drug similarity network.

### Enhanced efficacy and reduced toxicity in inter-cluster drug combinations on AML patient samples unveiled by cell viability drug screening

In the testing of all 16 inter-cluster and 12 intra-cluster combinations at five different concentrations, the cell viability of sixteen AML patient samples and five healthy donor samples were monitored. Patient samples with blast percentage more than 49% were chosen for testing with the Cell Titer-Glo (CTG) assay (Table S5). The average inhibition across dosages on 16 patient samples is regarded as efficacy, whereas the average inhibition across dosages on healthy samples is regarded as toxicity. The drug combinations with rectangular labels have higher efficacy and lower toxicity than the median. The proportion test (p-value *=* 0.006) revealed that the percentages of inter-cluster drug combinations with high efficacy (efficacy higher than the third quantile of efficacy values) and low toxicity (toxicities lower than the first quantile of toxicities) are significantly more than random choices.

The synergy and combination ratio (CR) of drug combinations on AML and healthy samples was then calculated using synergy scoring functions has ^25^, bliss ^26^, loewe ^27^, and ZIP ^28^ (Figure 4 and S3). The same analysis was done on synergy scoring values, and it was discovered that inter-cluster drug combinations differ significantly from random choices (P-values shown in the Figure 4 A-F). The drug combinations shown with rectangular labels have the highest synergy on AML patient samples, and the lowest synergy on healthy samples. Table 2 summarizes all six plots and the significant drug combinations according to different measures are highlighted by green (inter-cluster), yellow and purple (intra-clusters). Following CTG analysis, consensus across synergy scoring functions led to the selection of the five best drug combinations out of 28 to quantify blast-specific drug responses with flow cytometry.

**Fig. 4.**
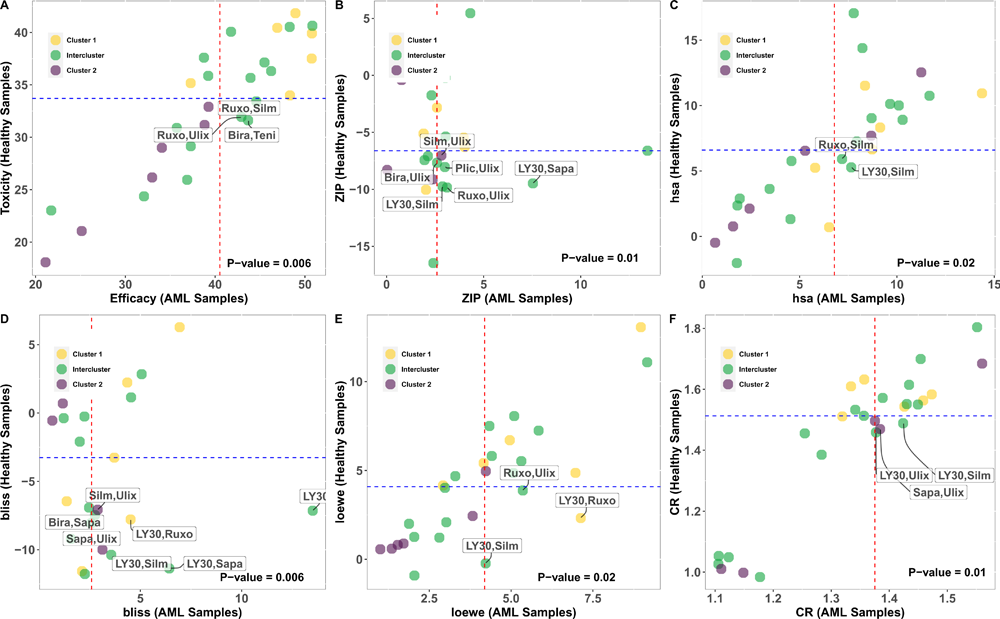
Drug combinations’ synergy scores on 16 AML samples and 5 healthy samples. X-axis depicts the synergy on AML samples and the Y-axis represents the synergy on healthy samples. The median inhibition on AML and healthy samples, respectively, is shown by the dashed lines in red and blue. There are three groupings: clusters 1, 2, and inter-cluster, and the color of each dot indicates each of these groups.

**Table 2.**
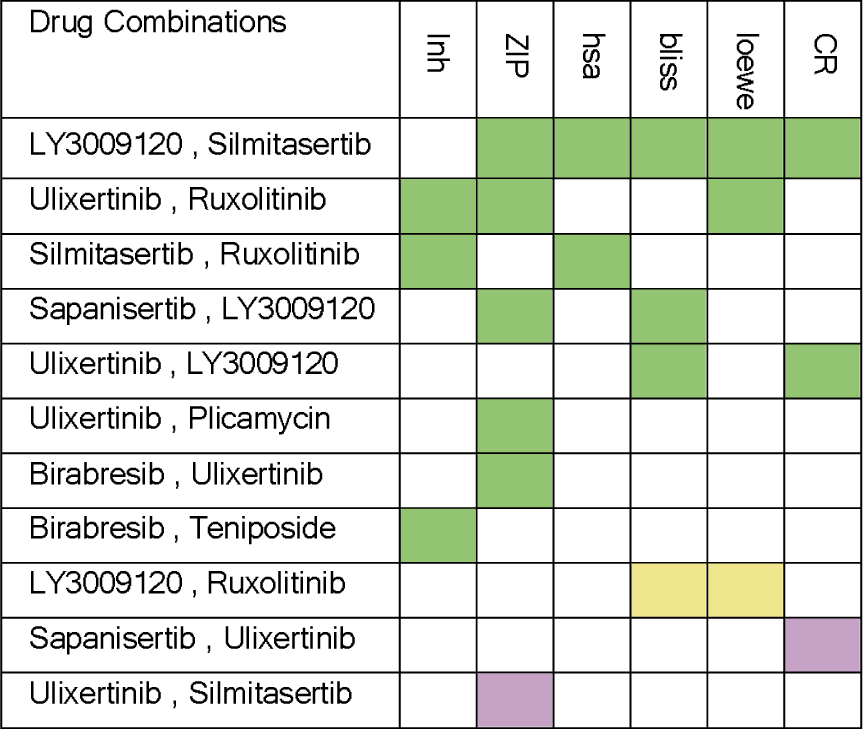
Selected drug combinations sorted by synergy scoring functions. Highlighted in green (inter-cluster), yellow (intra-cluster 1), and purple (intra-cluster 2). Inh stands for the average of inhibition of drug combinations on dosages, ZIP, hsa, bliss, and loewe are synergy scorings and CR is the combination ratio.

### Cell subtype viability analysis highlights low toxicity of selected combinations

Using the CTG assay, we measure the general bone-marrow mononuclear cell (BM-MNC) sensitivity, whereas with flow cytometry analysis we measure the number of live cells among different cell populations. Following 72-hour treatment with the 5 selected combinations on 3 different samples, viability of different cell subtypes of interest was measured by flow cytometry. For each sample, there is a specific plate layout which can be found in Figure S1. We used six cell surface markers (CD14, CD15, CD45, CD38, CD117, CD34) (Table S6) to identify the major leukocyte populations present in the AML BM-MNCs: monoblasts, myelocytes, leukemic blasts, leukemic stem cells, and myeloid progenitor cells (Figure S2).

In the studied samples, the average of blasts out of CD45 positive leukocytes, was 70% in DMSO, while on average 36% ± 16% of the blasts were killed by the combinations (Table S7). Based on the results, the percentage of dead cells for all 5 combinations in lymphocytes is considerably lower than 25% (Figure 5). More importantly selected combinations have lower synergistic effect on lymphocytes compared to the blast population, demonstrating the lower toxicity of combinations (Figure 7 and S3). The combination of JAK1/2 inhibitor (ruxolitinib) with either ERK or CSNK2A1 inhibitor has the highest efficacy and lowest toxicity which represents the important role of this target in AML. Numerous studies show the significance of the JAK/STAT signaling system in determining how hematopoietic cells react to various cytokines and growth factors ^29,30^. Recently there has been increased interest in different drug combinations with ruxolitinib ^31–34^ and as our results show the combinations of this drug, by having the lowest toxicity, seems to be promising for AML treatment.

**Fig. 5.**
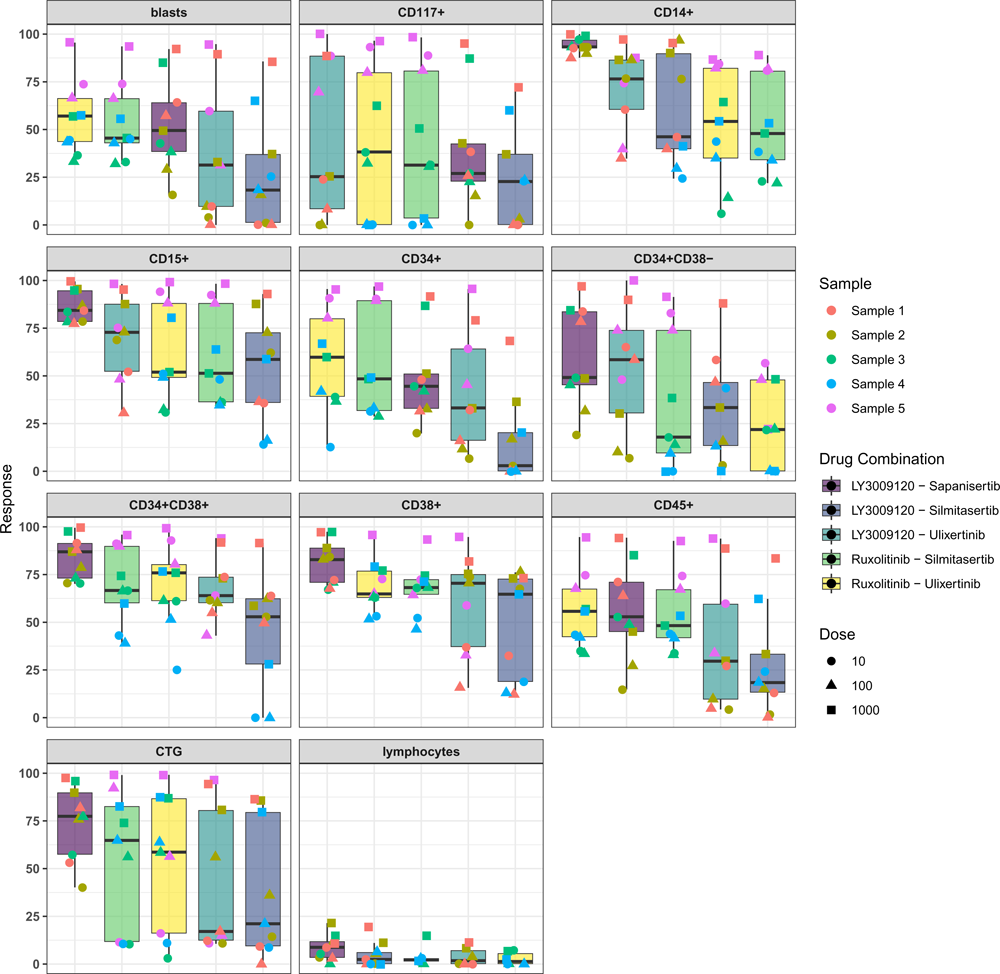
The cell viability assay (CTG) and response of different cell populations to 5 selected combinations using flow cytometry assay. Response signifies the percentage of dead cells following a 72-hour treatment. For each combination three different samples, distinguished by the color of points, were treated with three doses (10, 100, 1000), which are illustrated by different shapes of points. The colors in each cell group facet corresponds to a specific drug combination.

### Blast-specific drug responses in AML: Efficacy profiles of selected combinations

We were able to assess blast-specific drug combination responses and compare them to the other combinations within different samples. Among the five tested combinations, two combinations with ruxolitinib which targets JAK1/2 were among the most efficient combinations. The combination of ruxolitinib with ulixertinib, an ERK inhibitor, exhibits the strongest efficacy against blasts, according to the results. After treatment, the combination induced 47% ± 13% cell death in blasts (Figure 5 and Table S7) with a more synergistic effect on the blast population compared to the lymphocyte population (Figure 7A). We depicted the gating of 1000 nM concentration of each drug on sample AML_3 to better understand the impact of combination therapy vs. DMSO control and single drug treated samples in Figure 6. The number of blast cells in the ruxolitinib and ulixertinib treated well was reduced to 37%, showing the largest reduction compared to all other treatments, as shown in Figure 6A. The second combination of ruxolitinib and silmitasertib, a CSNK2A1 inhibitor, showed high efficacy on blasts. On average, this combination induced death to almost half ± 14% of the blast population but had less effect on lymphocytes (Figure 5 and Table S7). Additionally, this combination had a substantially higher inhibition rate compared to each single drug and acted synergistically towards the blast population (Figure 7A).

**Fig. 6.**
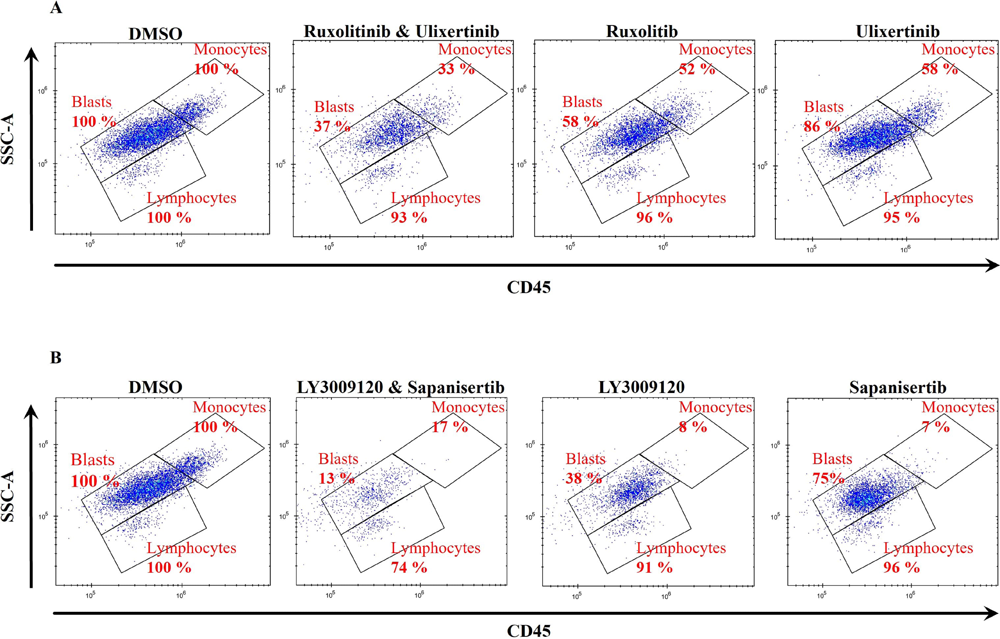
Flow cytometry scatter plots showing the effects of drug combinations on cell populations, along with comparisons to DMSO and single drug treated samples. (A) This figure illustrates the effects of ruxolitinib and ulixertinib combination and (B) LY3009120 and sapanisertib combination on blasts, monocytic cells (CD14+) and lymphocytes after 72h drug treatment. Numbers are presenting the percentage of cell counts in each cell population. The plot represents concentration of 1000 nM on sample AML_3.

**Fig. 7.**
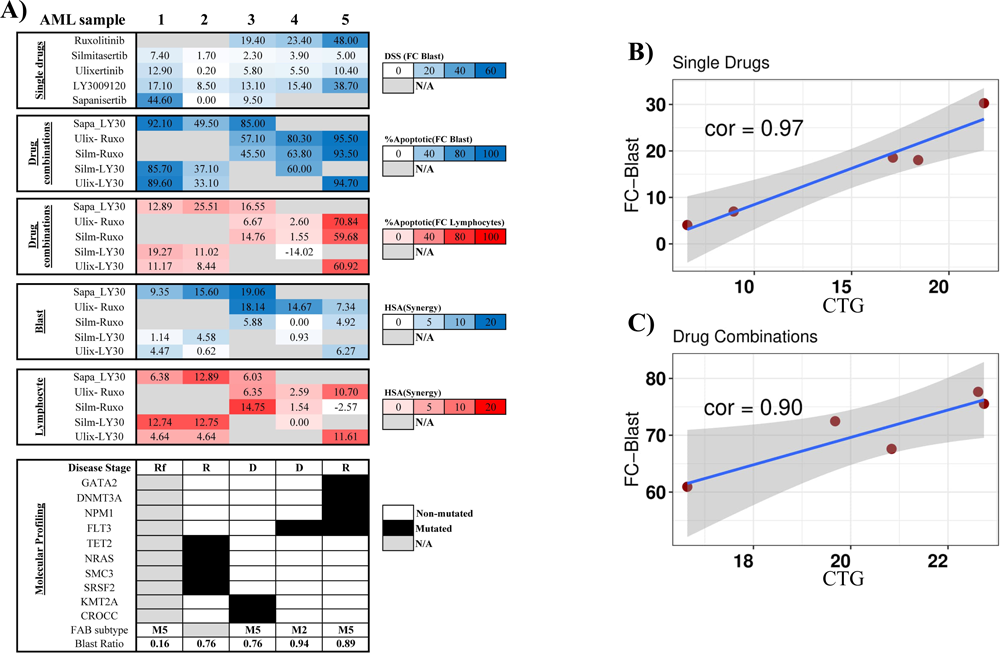
Characteristics of drug responses and correlation analysis in AML treatment: Comprehensive flow cytometry and CTG assessment. (A)Heat map showing characteristics of single agent and combinations response measured by flow cytometry readout. Blast-specific response of single drugs is highlighted according to drug sensitivity score (DSS) values with dark blue corresponding to high DSS value and white to low DSS value. Blast-specific and lymphocyte-specific response combinations in 1000 nM are highlighted according to percentage of apoptotic/dead cells, with dark blue in blast and red for lymphocyte corresponding to high percentage and white to a low percentage of apoptotic/dead cells. The synergistic effect of the drug combination was assessed based on the HSA synergistic score in 1000 nM on blast cells shown in blue and lymphocytes shown in red. (B) The correlation between responses measured by CTG and flow cytometry on five single drugs ruxolitinib, silmitasertib, ulixertinib, LY3009120, and sapanisertib, and (C) five drug combinations sapanisertib-LY3009120, ulixertinib-ruxolitinib, silmitasertib-ruxolitinib, silmitasertib-LY3009120, and ulixertinib-LY300912.

Given the importance of pan-RAF inhibition, we next examined LY3009120 in combination with three other drugs. The samples used for the combination of LY3009120 and sapanisertib (mTOR1/2 inhibitor), consist of 56% blast and the response for them is 40% ± 12% inhibition. To confirm that this combination is efficient, we analyzed the effect of LY3009120 and sapanisertib combination with single treated and DMSO treated cells in AML_3. In the combination treated sample, the blast cells were significantly reduced to 13% while in the individual drugs LY3009120 and sapanisertib reduced the blasts to 38% and 75%, respectively (Figure 6B). These results indicate that this combination has substantially higher inhibition rate compared to each single drug and a greater synergistic effect on blasts than on lymphocytes (Figure 7A). Ulixertinib (ERK inhibitor) is the second drug that is being used in combination with pan-RAF inhibitor. Patient samples treated with this combination, on average, contain 60% blast cells and after treatment they are reduced to 28% ± 14%. Finally, we tested the combination of LY3009120 with silmitasertib, a CSNK2A1 inhibitor on three different samples. The average blast population for these three samples is 62% and the response was 21% ± 5%. Overall, as shown in Figure 5 and 7A, all combinations have very little impact on the lymphocyte populations, demonstrating low toxicity, and significantly more impact on less differentiated malignant cells, demonstrating the efficacy of the combinations.

### Increased sensitivity of AML samples to combination therapies over single drugs, regardless of genetic mutations and prognosis categories

There is a significant correlation between CTG assay and blast specific results, indicating that reduction in cell number measured by CTG, is related to the malignant cell populations (Figure 7B and 7C). The cell viability readout for a single drug is converted to a drug sensitivity score (DSS) which is a drug sensitivity metric based on area under the dose-response curve. A greater DSS indicates higher sensitivity^35^. Strikingly, by combining selected inter-cluster drugs, the blasts were targeted, and combinations showed a synergistic effect on this population (Figure 7A). Then in order to monitor the drug responses based on genetic changes, we examined the existing mutations in selected samples that are the most significant mutations among AML patients (Figure 7A) ^36,37^. To evaluate the impact of the combinations on samples bearing genetic alterations, some mutations that are frequently found in AML patients were considered (Figure 7A). *FLT3* mutation is a well-known mutant gene among AML patients and is represented in two samples. The other prevalent mutations were *NPM1, GATA2, DNMT3A, TET2, KMT2A, NRAS, SMC3* and *SRSF2*. The combinations induced a synergistic effect on the blast population, regardless of the genetic alterations. The European Leukemia Network (ELN) classifies patients into three prognosis categories: “favorable”, “intermediate”, or “adverse”^38^. AML patients are also classified based on morphological features which is named French-American-British Classification (FAB) ^39^. Regardless of sample type, we observed a synergistic effect following treatment. Importantly, after therapy, we noticed a synergistic effect in all samples, indicating that these combinations are effective at combating the heterogeneity of the AML disease. It has been demonstrated that drugs should target the less differentiated leukemic blasts to achieve the best response in patients ^6^. Given these two facts—the presence of the most relevant mutations and the prevalence of blast cells in the samples— the combinations seem to be promising for treatment.

### Efficacy and toxicity of the novel combinations compared to first-line treatment in AML

In the following analysis, we compared the proposed combinations in this study (ruxolitinib-ulixertinib and LY3009120-sapanisertib) with two FDA-approved combinations (venetoclax-azacitidine and venetoclax-cytarabine), as well as the investigational combination of venetoclax-ruxolitinib. As illustrated in Figure 8, venetoclax-ruxolitinib demonstrates the highest efficacy on both blast cells and lymphocytes compared to the other combinations. This dual efficacy profile is a noteworthy advantage; however, it comes at the cost of heightened toxicity, as indicated by our results.

**Fig. 8.**
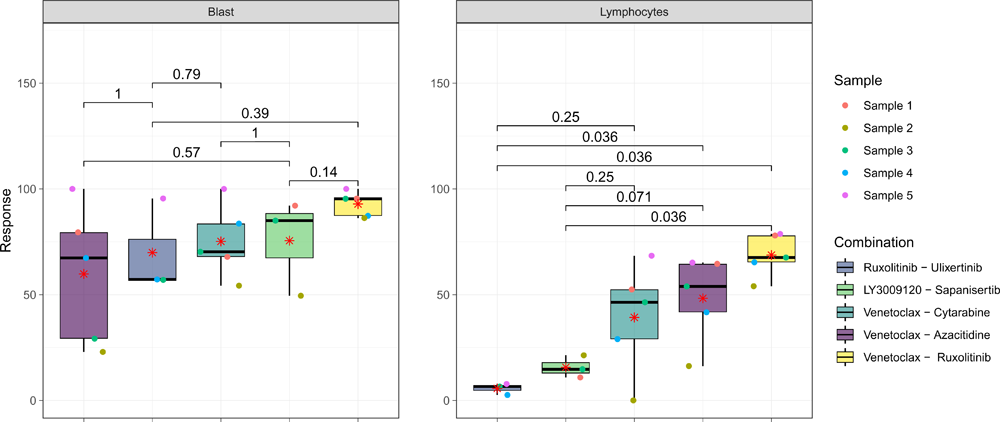
Flow cytometry assay of selected combinations compared to first-line AML combinations. Response signifies the percentage of dead cells following 72-hour treatment. Each combination has been tested on different samples at 50 nM concentration for venetoclax and 1000 nM for the other drugs. Red asterisks define the average response for each combination and colored dots represent different samples. Each panel also represents six p-values resulting from the Wilcoxon signed-rank tests to compare statistically two proposed combinations with three other combinations (including two first-line treatments and one investigational combination) for AML.

Conversely, the novel combinations, ruxolitinib-ulixertinib and LY3009120-sapanisertib, show comparable efficacy in targeting blast populations as the established combinations. Notably, there is no significant difference in terms of efficacy (p-values are shown in Figure 8). However, these two combinations have a significant advantage in demonstrating lower toxicity compared to first-line combinations, particularly for lymphocytes. The effect of ruxolitinib-ulixertinib and LY3009120-sapanisertib on blast lymphocyte population is significantly lower than all other combinations except for venetoclax-cytarabine (p-value = 0.25) which is not significant but still is lower. This reduction in toxicity suggests these combinations can offer effective treatment while minimizing side effects associated with prevalence therapies.

## Discussion

In this study we employed a nominal data mining approach to construct a weighted bipartite network for the selection of the most effective ^10^, as well as the least toxic drugs. The selected combinations inhibit important signaling pathways and included pan-RAF inhibitors, JAK1/2 inhibitors, BET family inhibitors, topoisomerase II inhibitor, CSNK2A1 inhibitors, ERK inhibitors, mTOR1/2 inhibitors, and DNA binding drugs.

The MAPK (RAS/RAF/MEK/ERK) signaling pathway is hyperactivated in AML patients, leading to leukemogenesis, leukemia progression, and chemo resistance ^40–43^. Since, targeting RAS and ERK remains challenging, pan-RAF inhibitors are an interesting new pharmacological class in this context ^44,45^. Recent studies demonstrated that the pan-RAF inhibitor LY3009120 induces growth inhibition and apoptosis in RAS-mutated AML cell lines ^46,47^. Another component of the MAPK pathway is ERK, and the ERK inhibitor ulixertinib is a promising early efficacy agent for treating patients with tumors harboring alterations in the MAPK pathway ^48^. The ERK pathway is activated in approximately 30% of all human cancers via *RAS, BRAF,* or *MAP2K1 (MEK1)* activating mutations and thus targeted therapeutics are vital to treating ERK pathway–activated human cancers ^49^. Bhagwat et al. demonstrated the efficacy of LY3214996 as an ERK inhibitor to delay or reverse resistance to BRAF and MEK inhibitor therapy. They also demonstrated a synergistic effect of the combination of LY3214996 with a pan-RAF inhibitor, LY3001920, in a KRAS-mutant colorectal cancer xenograft model ^50^. Moreover, here we identified ulixertinib and LY3009120 as an inter-cluster combination with high efficacy and low toxicity. Upon further analysis, we found two combinations of ulixertinib and three combinations of LY3009120 among the five most efficient and synergistic combinations.

The other signaling pathway is JAK signaling, which plays critical roles in several important intracellular signaling pathways ^51,52^. In a number of leukemias, aberrations of the JAK/STAT pathway and constitutive STAT activation have been well described ^53^. Ruxolitinib has been approved as a non-selective JAK1/2 inhibitor for patients with intermediate to high risk primary or secondary myelofibrosis ^54^. Ruxolitinib, by inhibiting JAK 1/2, reduce activation of JAK-signal transducers and lower down the transcription (STAT) signaling ^55^. Based on our computational analysis, CTG results, and flow cytometry analysis, we revealed that combinations of ruxolitinib with ulixertinib and silmitasertib are the most blast-specific compounds, while having minor effect on lymphocytes in AML *ex vivo* screening.

Next, we examined the effect of Bromodomain and Extra-Terminal motif (BET) protein inhibitors on cancer cells as BET proteins are often altered in cancer. While bromodomain inhibition is effective, not all patients benefit from it and become resistant over time ^56^. Researchers recently reported the efficacy of combined JAK/BET inhibition on cytokine production, BM fibrosis, and tumor burden *in vivo* ^57^, leading to complete reversal of reticulin fibrosis in mice. In our model, this drug was classified among effective drugs in cluster 1 and based on the CTG assay, in combination with sapanisertib, ulixertinib, and teniposide can be potential compound for the treatment of AML patients.

The other drug that was selected is teniposide which is a chemotherapy medicine used to treat solid tumors and leukemia, typically in combination with other antineoplastic agents. This agent causes DNA damage in proliferating cells by inhibiting topoisomerase II. Recently, it has been proven that teniposide not only inhibits topoisomerase II but also inhibits MYB, transcription factor, activity and expression ^58^. Topoisomerase II inhibitors’ ever-evolving role in the treatment of AML has led to an increase in their utilization in combination therapies ^59^. Here in this study, we identified the combination of teniposide and birabresib among the most effective compounds with low toxicity.

The next drug is silmitasertib that inhibits CSNK2A1. Studies showed increased expression of this kinase in AML cells, which contributes to several biological processes related to cancer progression such as apoptosis, cell growth, proliferation and transcription ^60^. Silmitasertib is currently being evaluated in clinical trials for the treatment of multiple malignancies ^61^. Hereby, considering the importance of pan-RAF and CSNK2A1 inhibitors, targeting these two pathways simultaneously might be a promising treatment. Additionally, as we suggested, the combination of silmitasertib and ruxolitinib could be considered as potential combinations for treating AML patients.

We then, studied the effect of plicamycin, which inhibits RNA synthesis by binding to GC-rich regions of DNA. Plicamycin has potent anti-tumor effects, both as mono-therapy and in combination with anti-tumor drugs ^62^. A highly selective adenosine triphosphate-competitive inhibitor of mTOR kinase, named sapanisertib has been also included in this study. Sapanisertib has demonstrated antitumor activity in AML by inhibiting both mTORC1 and mTORC2 ^63^. Sapanisertib in combination with LY3009120, birabresib or ulixertinib has synergistic effect in AML samples (Figure 4).

Lastly, we thoroughly compared the two FDA-approved combinations (venetoclax-azacitidine and venetoclax-cytarabine) and one investigational combination (venetoclax-ruxolitinib) with the novel combinations suggested in this study, namely ruxolitinib-ulixertinib and LY3009120-sapanisertib. Notably, the outcomes validated the comparability of the suggested combinations’ efficacy with first-line combinations in this investigation. There is no significant difference in efficacy on blast cells, between proposed combinations and first-line AML combinations. Significantly, the toxicity of selected combinations is lower than others, except for venetoclax-cytarabine, which indicates that proposed combinations might offer both effective treatment and a reduced side effect compared to standard AML combinations.

In summary, we proposed effective drug combinations for AML patients with the highest efficacy and lowest toxicity based on nominal data mining method and *ex vivo* drug sensitivity assay. The combinations were tested on three different samples, and the results indicate that the combinations of ruxolitinib-ulixertinib and sapanisertib-LY3009120 have the highest synergistic effect, the highest efficacy on blasts, and the lowest toxicity. Although the approach of combining targeted agents suffers from cumulative toxicity effects ^64^, we have demonstrated that our approach overcomes this limitation in designing a drug combination in AML. Considering the importance of toxicity, in all steps we considered toxicity as an important factor for the selection of combinations with the lowest effect on healthy cells. Standard chemotherapy kills most of the blasts as well as other cell types and has a high value of toxicity ^6^, while recommended combinations in this study are effective on blasts, but have lower toxicity on other cell populations.

## Acknowledgments

We thank the patients and donors who participated in the study. Drug screening was carried out at the FIMM High Throughput Biomedicine Unit (HTB), which is hosted by the University of Helsinki and supported by HiLIFE and Biocenter Finland. Alun Parsons is gratefully acknowledged for his technical support. We also appreciate comments from Marc Baumann, Julio Saez-Rodriguez and Farnaz Barneh. This study was financially supported by the Research Council in Finland [grant 332454 to M.J.; grants 320185, 334781, 357686 to C.A.H.], and Jane & Aatos Erkko Foundation [grant 220031 to M.J.], and the Sigrid Jusélius Foundation [grant to C.A.H.].

## Authorship Contributions

M.J. conceived of the study and designed research questions. M.J. and C.A.H. oversaw the scientific direction, supervised the project, and assisted with the manuscript. Also C.A.H. contributed resources including patient samples. M.M. led the computational and statistical analysis, interpreted the data, and assisted with study design, while E.G., T.R., and J.S. designed and developed the experimental methods for cell viability analysis. J.M. and F.I. contributed to the interpretation of the findings for drug sensitivity analysis. M.M and S.M contributed bone marrow samples from healthy donors.

## Data availability statement

The data that support the findings of this study and all code for data analysis are openly available at https://github.com/jafarilab/DrugComb_AML.

