## Supplementary data 1 for "Designing patient-oriented combination therapies for acute myeloid leukemia based on efficacy/toxicity integration and bipartite network modeling"

Figures

Tables

### 1. Materials and Methods

*Ex vivo* drug-response data were generated at the Institute for Molecular Medicine Finland (FIMM) for a prospective series of 252 samples from 186 patients with acute myeloid leukemia (AML)^1^. The results of experiments using the dead cells (CellTox Green) readout were extracted, which included the drug response of 625 chemical compounds on 199 bone marrow AML samples from patients. To determine the inhibition efficacy of each drug on each sample, the mean value of drug response across dosages was extracted after processing. This dataset can be represented as a 199-by-624 matrix, with rows representing samples and columns representing drugs tested on AML samples. Furthermore, the full submatrix (with no missing entries) of 81 patient samples and 296 chemical compounds was extracted using the NIMMA package^2^. The magnitudes of the dose response levels vary across experimental protocols and techniques due to the heterogeneity of the different platforms on which the high-throughput assays were performed. We normalized the mean value of dose response levels to provide coincident and comparable therapeutic efficacy across different experiments to facilitate downstream use of our dataset. Given that cell death was used in drug sensitivity assays, we calculated the inhibition rate (*R_inhibition_*) of cancer cells to drug treatments as a uniform measure using the min–max normalization method:

$$R_{inhibition}=\frac{celldeath-min(celldeath)}{\max\left( celldeath \right)-min(celldeath)}$$

As a result, one represents the highest sensitivity, and zero represents the lowest sensitivity, since the normalized inhibition rates range from 0 to 1. In matrix *A* each *ij* entry denoted by *a_ij_* indicates the normalized inhibition rate of drug response *j* on sample *i*.

1.1. Reconstruction and analysis of the bipartite network model. A weighted network *G* = (*V, E, ω*) is a triple—a set of three elements—in which *V* is a set of nodes, *E* is a set of edges between nodes in *V*, and *ω* is a function that assigns a weight to each edge $e\in E$. A network is said to be bipartite if *V* can be divided into two sets, *V*_1_, *V*_2_, so that every edge $e\in E$ is connected to a node in *V*_1_ and a node in *V*_2_. A bipartite weighted network is shown as *G* = (*V*_1_*, V*_2_*, E, ω*). Suppose $S= \left\{ s_{1},s_{2},\ldots,s_{m} \right\}$ and $D= \left\{ d_{1},d_{2},\ldots,d_{mn} \right\}$ are samples and the drugs sets in the dataset, respectively. The data matrix *A* was used to construct a weighted bipartite network where *V*_1_ = *S* was set of 81 samples, and *V*_2_ = *D* consisted of 296 drugs. The weight of the edge that joins node *s_i_* (sample *i*) and node *d_j_* (drug *j*) was the *ij* entry of the matrix *A*. A weighted bipartite network was built, with two parts: samples and compounds, and weight representing the inhibition rate (*R_inhibition_*) as explained above.

1.2. Construction and analysis of the drug similarity network. A bipartite network can be projected into two different types of unipartite networks containing nodes of only one type. The projection of the bipartite network, A, onto the "drug" node set was considered here, and the weight of edge between drug *d_i_* and drug *d_j_* was as follows:

$$w_{ij}= \sum_{k=1}^{81} (a_{ik}\times a_{jk})$$

This weight was considered as the similarity score between two drugs, *d_i_* and drug *d_j_* , according to their efficacy on samples. Only edges with a weight greater than the median of similarities were kept in order to consider them strong enough edges in the projected network. In order to identify functionally similar drugs in terms of drug response the Louvain community detection method^3^ was used.

1.3. Computational corroboration. To accomplish intra-cluster homogeneity analysis, we employed two computational methods. The first method identified the significant difference between biological pathways of drug targets’ protein targets at each cluster, while the second evaluated drug chemical structure similarity at each cluster. Using the drug-target common (DTC) database^4^, we built a drug-target network, which was a bipartite network in which each link connects drugs to their protein targets.

To better understand the protein targets of drugs in each cluster, we assign a score to each protein based on the number of distinct drugs targeting that protein in clusters 1 and 2. Let *f*_1_*_,P_* (*f*_2_*_,P_* ) denotes the number of unique drugs in cluster *C*_1_ (*C*_2_) targeting a particular protein P. The score of protein P, defined by

$$S\left( P \right)= \log\frac{f_{1,P}}{f_{2,p}}$$

Proteins with a score S greater than log (2) are considered to be preferentially targeted by drugs in cluster 1, denoted by PPT1. Similarly, PPT2 proteins have a score of less than *log*(0*.*50). The KEGG pathway annotations and biological processes of each cluster’s protein targets were also extracted using clusterprofiler R package^5^ and ShinyGO^6^. The KEGG pathway annotations and biological processes provided in the package were used to map pathways and biological processes (GO) to our protein sets PPT1 and PPT2. The settings used in the gseKEGG and gseGO functions were 10,000 permutations, the minimum size of the gene set to test was 10, and the maximum size of the gene set to test was 500. REVIGO was used to summarize the enriched GO terms [(http://revigo.irb.hr/)](http://revigo.irb.hr/)). The significantly enriched GO terms (Adj.Pvalue < 0.05) were analyzed by REVIGO^7^. This program removes redundant GO terms and the similarity between terms is reflected by semantic space.

A simplified molecular input line entry system (SMILES) of the drug molecules was retrieved in order to compare the chemical structures of the compounds, and it was then converted into an extended connectivity fingerprint (ECFP) in order to evaluate the dice similarity between the molecules. The dice similarity between molecules A and B is one of the standard metrics for molecular similarity calculations in which

*S_A,B_* = 2*c/*(*a* + *b*)*,*

where a is the number of ON bits in molecule A, b is the number of ON bits in molecule B, and c is the number of ON bits in both A and B molecules^8^. To calculate the dice similarity of the compounds, a simplified molecular input line entry system (SMILES) of the drug molecules was retrieved and transformed into an extended connectivity fingerprint (ECFP). The rcdk package^9^ was used to calculate the similarity between chemical compounds.^10-12^

We utilised four well-known scoring functions ZIP^13^, HSA^10^, Bliss^11^, and loewe^12^ to assess the potential synergy of drug combinations. The observed drug combination responses in these models were compared with the expected combination responses to quantify synergy of drug combination. The combination ratio (CR) was also defined as the ratio of the response of combinations to the maximum for the two single agents, respectively. By this metric, a CR value of higher than 1 indicates the drug combination is more effective than either single agent^14^. The effect of drug combinations on five dosages (1,10,100,1000,10000) was monitored in this study, and the DECREASE model was used to predict drug combination dose-response at the full matrix. Synergy scores were calculated using the SynergyFinder web application (version 3.0)^15^.

1.4. Patient samples. Freshly frozen bone marrow mononuclear cells (BMMC) from 16 AML patients and 5 healthy volunteers were obtained from the Helsinki University Hospital Comprehensive Cancer Center after informed consent (permit numbers 303/13/03/01/2011, Helsinki University Hospital Ethics Committee). The samples were numbered from 1 to 16 in supplementary Table S4, from which samples 1 to 5 have been used for both CTG and FC analysis. The samples were selected based on clinical malignant cells of higher than 49%. Following thawing, the cells were cultured in RPMI supplemented with 12.5% HS-5 stromal cell derived conditioned medium (CM), 10% fetal bovine serum, 2mM L-glutamine and penicillin/streptomycin and DNAse, then incubated at 37°C and 5% CO2 for 2-3hours. After the incubation time the cells were counted and adjusted to a final concentration of 200,000 and 1 106 cell/ml for CTG and FC, respectively. The patient characteristics were presented in the Online Supplementary Table S1.

1.5. Preparation of drug plates. The compounds (Supplementary Table S2) were dissolved in dimethyl sulfoxide (DMSO) and dispensed on 384-well plates (Corning, Corning, NY, USA) using an acoustic liquid handling device Echo 550 (Labcyte, Sunnyvale, CA). DMSO was used as negative control and 100 µM benzethonium chloride (BzCl) as positive control (Table S2).

1.6. Cell viability analysis using Cell Titer-Glo. The AML cells were seeded on pre-drugged 384-well plates (Corning) containing chemical compounds at five different concentrations in two replicates. The final volume of cell in each well was adjusted to 5000 cells in 25 µl per well and incubated for 72h at 37°C and 5% CO2. Cell viabilities were assessed using the CellTiter-Glo 155 assay (Promega), and the luminescence signal was measured using a PHERAstar FS plate reader (BMG LABTECH, Ortenberg, Germany). As quality control, viability screening was used to check how the cells survive in 384-well plates during the 72h incubation. Viability of the cells was monitored at 0h and at 72h using the CTG assay.

**1.7.** High throughput flow cytometry. For the phenotype-based drug sensitivity profiling, the high throughput flow cytometry (HTFC) assay was performed. Following thawing BMMC were seeded using MultiFlow FX.RAD (BioTek) to 384-well compound plates (Greiner), 20,000 live cells in 20 µl CM in each well, and incubated for 72h at 37°C and 5% CO2 (Figure S1). Monoclonal antibodies CD45, CD38, CD34, CD117, CD11b, CD14 and CD15, apoptosis dye Annexin-V and dead cell exclusion dye DRAQ7 were added with Echo 525 (Labcyte Inc.) and stained for 30 min at room temperature (Table S3). Cells were analyzed with iQue3 (Sartorius, Germany) HTFC. ForeCyt software (Sartorius) was used to analyze the remaining viable cells and normalized to the number in the DMSO control wells. Drug sensitivity scores (DSS) and SynergyFinder 2.0 were used to analyze the results (references). The gating strategy is presented in Figure S2.

1.8. Statistical Analysis. T-test was used to show that the mean of inter-cluster dice similarities is less than the mean of intra-cluster similarities. We also used a statistical proportion test to show that the proportion of inter-cluster drug combinations with efficacy greater than the third quantile (*Q*_3_ or 75th percentile) of efficacy values and toxicity less than the first quantile (*Q*_1_ or 25th percentile) of toxicity values is significantly higher than the random choices (*probability* = 0*.*33). This demonstrates that inter-cluster drug combinations have the highest efficacy and the lowest toxicity. The similar approach was utilized for calculating CR values as well as synergy scores. In KEGG, a biological pathway enrichment analysis was calculated based on hypergeometric test followed by false discovery rate (FDR) correction. Fold Enrichment was calculated by dividing the percentage of genes in the list that belong to a pathway by the corresponding percentage in the background. Fold Enrichment indicates how significantly genes from a specific pathway are over-represented^6^.

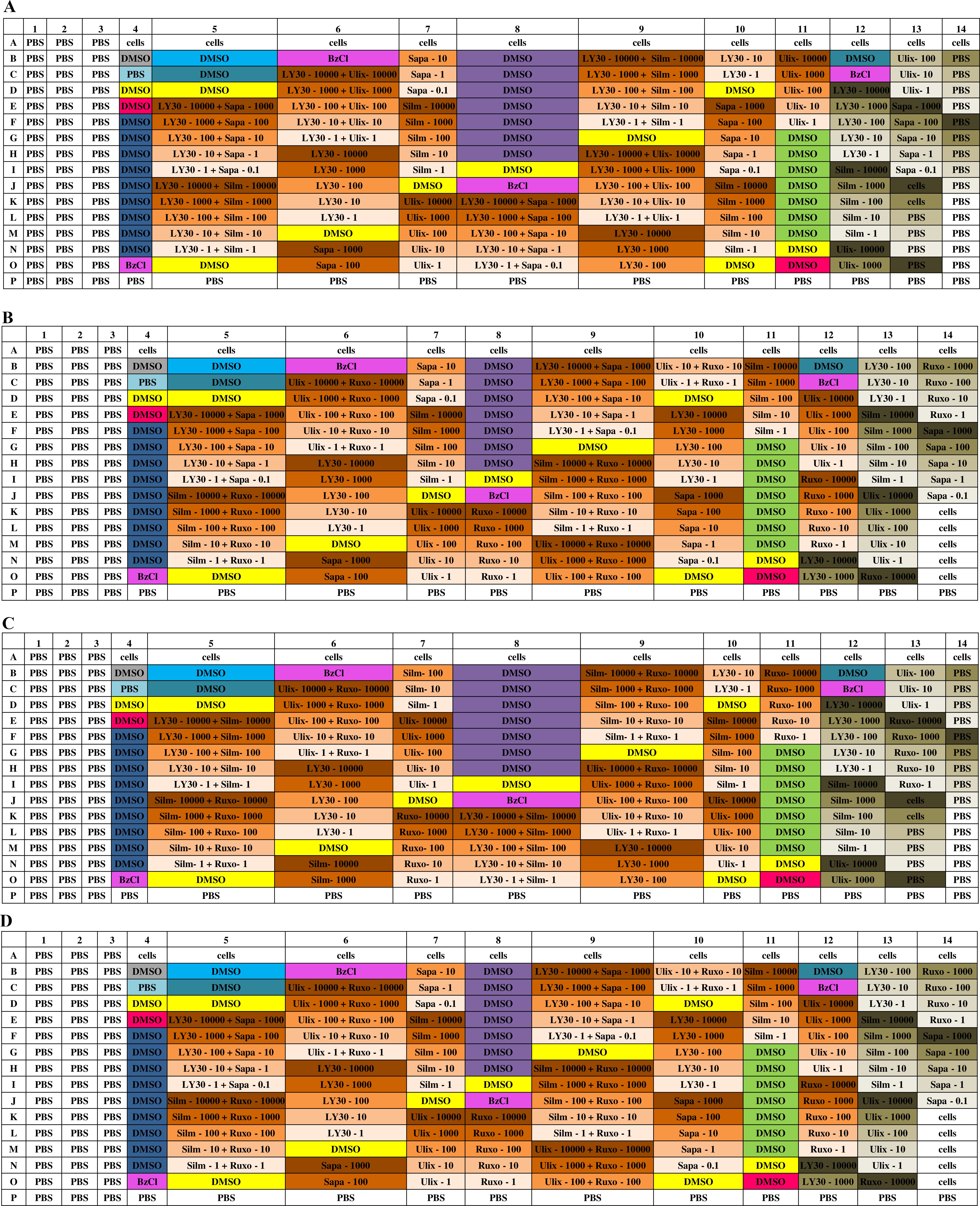

**Figure 1. Plate layout for FC assay on patient samples.**

This figure represents the 384-well plate drug design for sample (A)AML_1 and AML_2, (B)AML_3, (C) AML_4 and (D) AML_5. These layouts contain two replicates for each compound with 5 different concentrations, sapanisertib: 0.1, 1, 10, 100, and 1000 nM, and all other drugs: 1, 10, 100, 1000, and 10000 nM. LY3009120 (LY30), Teniposide (Teni), Ruxolitinib (Ruxo), Silmitasertib (Silm), Birabresib (Bira), Ulixertinib (Ulix), Plicamycin (Plic), Sapanisertib (Sapa).

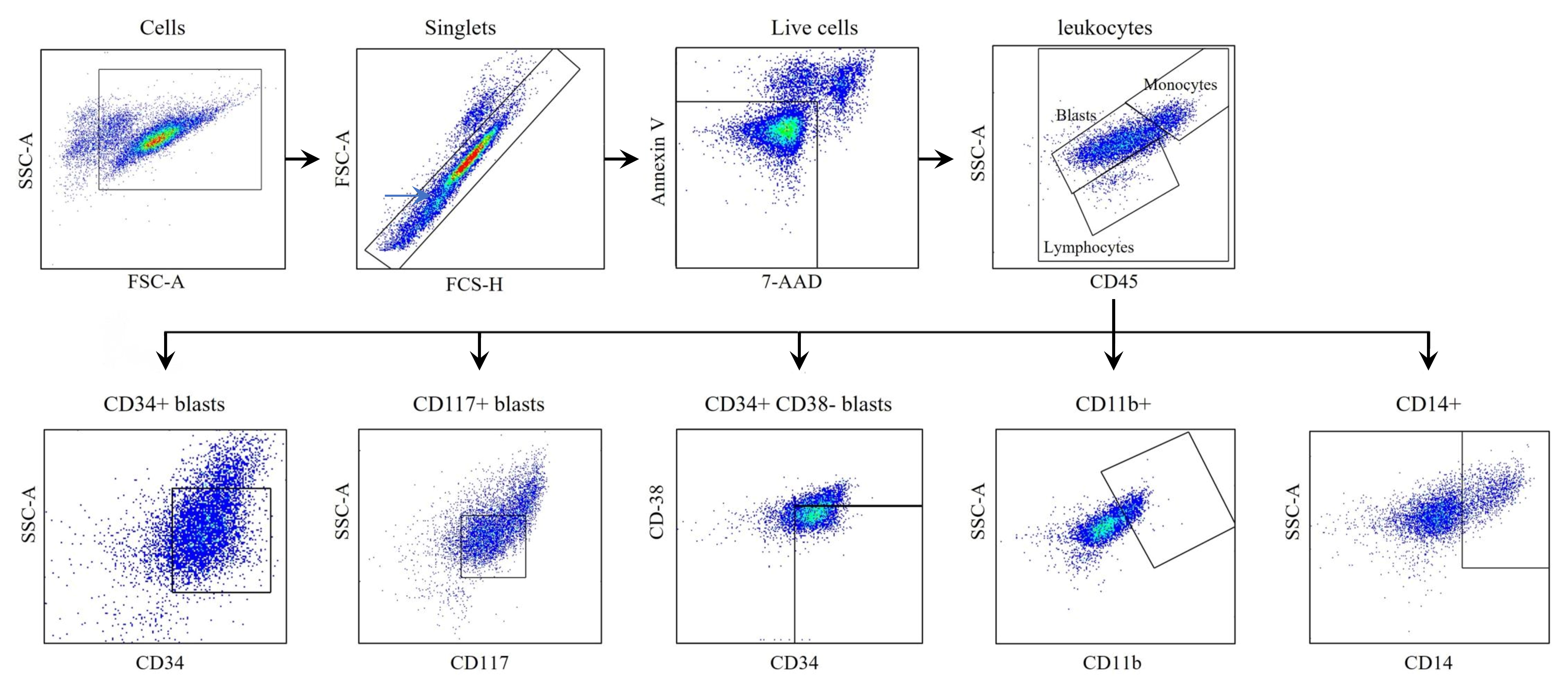

**Figure 2. Gating strategy of cell populations.**

Cells were gated and debris removed based on SSC-A/FSC-A after singlets were identified with FSC-A/FSC-H. Viable cells were gated by excluding the apoptotic and dead cells using Annexin V and DRAQ7, respectively. SSC-A/CD45 was used to gain an overview from the cell composition of the AML sample: blasts – SSC^low^/CD45^dim^, lymphocytes – SSC^low^/CD45^bright^, and monocytic cells – SSC^mid^/CD45^bright^. Blasts were identified with CD34 and CD117 antibodies and leukemic stem cells were gated as CD34+/CD38-. Additional markers CD14 and CD11 were used to gate cells differentiated towards monocytic and granulocytic lineages.

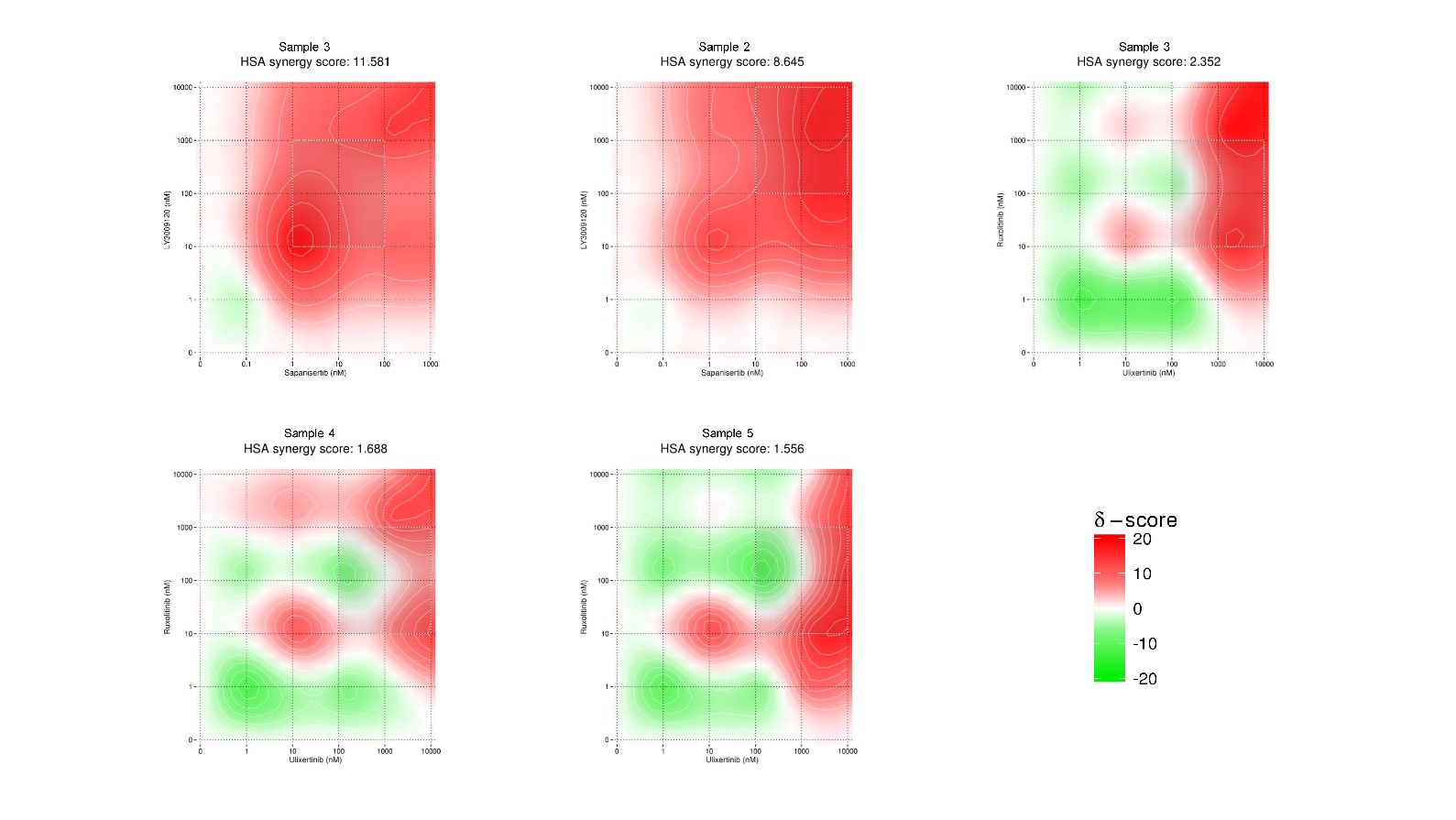

**Figure 3. Synergy analysis across dosage levels.**

This figure presents the synergy analysis results across a range of dosage levels for the reported drug combinations in AML blast populations. Each combination's synergy values are depicted at various dosage points, offering a comprehensive view of the impact of dosage on synergistic responses. The y-axis and x-axis illustrate the dosage levels for each drug, and synergy is depicted as color. Red color represents higher synergy and green represents lower synergy level.

**Table 1. List of drugs in each cluster**

| Cluster 1 | SGI-1776; Apatinib; Cediranib; Dabrafenib; Encorafenib; Icotinib; Lovastatin; Neflamapimod; Talmapimod; Tandutinib; Pravastatin; 1-methyl-D-tryptophan; Brivanib; Linifanib; Perifosine; Dovitinib; Fludarabine; Quisinostat; Valrubicin; MK-8745; VE-821; 8-chloro-adenosine; PF-4800567; AZD-8186; Alectinib; Capmatinib; Golvatinib; Sapitinib; TAK-285; Tesevatinib; Tucatinib; Vandetanib; BMS-777607; Roxadustat; EPZ-5687; GSK-J4; GSK2801; Lomeguatrib; Mocetinostat; Sepantronium bromide; Barasertib; Ibrutinib; Pemetrexed; AZ191; Tazemetostat; AZD4547; Ruboxistaurin; Infigratinib; Methotrexate; Birabresib; CEP-37440; GSK2656157; Marimastat; Verdinexor; Vismodegib; Tamoxifen; GDC-0919; Orteronel; Tasquinimod; BGB324; Galiellalactone; VER 155008; Everolimus; Vorinostat; Floxuridine; Pentostatin; Temozolomide; Allopurinol; Pomalidomide; Goserelin; Letrozole; Thalidomide; Enzastaurin; Lapatinib; Panobinostat; Valproic acid; ENMD-2076; Megestrol acetate; Rabusertib; Metformin; Plerixafor; Pazopanib; Losmapimod; Epacadostat; GSK2879552; Tacrolimus; Pracinostat; Cisplatin; Flutamide; Oxaliplatin; PH-797804; Rocilinostat; URB597; KU-60019; PHA 408; UCN-01; Vatalanib; AZD7545; GNE-7915; Tideglusib; Omipalisib; Tepotinib; Plicamycin; Tretinoin; Oprozomib; Vistusertib; Mubritinib; NMS-873; Pinometostat; Entinostat; Mitotane; Alvocidib; CUDC-305; BIIB021; Onalespib; BMS-911543; Chloroquine; Toremifene; LY3009120; Fingolimod; Doxorubicin; Aldoxorubicin; Ganetespib; IOX-2; Spebrutinib; TEW-7197; Bortezomib; Triciribine; Resminostat; Cladribine; Bimatoprost; Momelotinib; Ponatinib; Cilengitide; NVP-BGT226; Crenolanib; Ruxolitinib; PF-670462; UNC2881; SAR405838; Omacetaxine; Dexamethasone; Givinostat; PF-06463922; PCI-34051; GSK923295; CPI-613; Anastrozole; AR-42; Cerdulatinib; 8-amino-adenosine; AZD-5438; Afuresertib; Bosutinib; Enzalutamide |
| --- | --- |
| Cluster 2 | Hydroxyfasudil; Cobimetinib; Trametinib; AZD-5363; Bentamapimod; GSK-690693; RAF-265; TAK-901; Tivantinib; Ralimetinib; Regorafenib; Nilotinib; Erlotinib; Imatinib; Midostaurin; Vemurafenib; Crizotinib; Doramapimod; Masitinib; Seliciclib; Palbociclib; Neratinib; Pictilisib; Teniposide; SCH772984; PS-1145; Alpelisib; GSK-1070916; Lucitanib; Palomid-529; PF-00562271; Telatinib; Quizartinib; Sunitinib; GDC-0623; C646; GSK343; RGFP966; SGC-CBP30; SGC0946; Tubacin; Tubastatin A; Amuvatinib; Tamatinib; AZD1480; Cabozantinib; Fostamatinib; PF-04708671; Arsenic(III) oxide; Mirdametinib; AVN944; TH588; AT-406; PAC-1; Taselisib; FRAX486; Molibresib; AMG-232; PTC-209; UM729; MST-312; Glasdegib; GSK2830371; Varespladib; NVP-LGK974; Temsirolimus; Paclitaxel; Cytarabine; Rucaparib; Ixabepilone; Thioguanine; Azacitidine; Mitomycin; Methylprednisolone; Lenalidomide; Sonolisib; Triapine; Rociletinib; Fulvestrant; Ulixertinib; Ridaforolimus; Sirolimus; TGX-221; Axitinib; Carboplatin; WEHI-539; Foretinib; Dasatinib; Indibulin; GSK650394; Atorvastatin; Sapanisertib; APR-246; Gemcitabine; Saracatinib; AZ 3146; ML323; Itraconazole; LY-2874455; StemRegenin 1; Gedatolisib; Bafetinib; Celecoxib; Motesanib; Birinapant; Buparlisib; Filanesib; Volasertib; GSK269962; BI 2536; MK-2206; Abemaciclib; Copanlisib; Binimetinib; BMS-754807; Raloxifene; Nilutamide; Bleomycin; Simvastatin; Gefitinib; Baricitinib; Tosedostat; Tarenflurbil; Olaparib; TRAM-34; Sotrastaurin; Ipatasertib; Aminoglutethimide; GSK-2334470; Tacedinaline; Sonidegib; AZD7762; Silmitasertib; CEP-32496; Hydroxyurea; ASP3026; Veliparib; Docetaxel; Linsitinib; Niraparib; Uprosertib |

**Table 2. Biological processes**

| **Description** | **protein_group** | **weight** | **protein_list** |
| --- | --- | --- | --- |
| chromatin organization | PPT2 | 25 | Q9Y6K1, Q9UMN6, Q9H5I1, Q8WTS6, Q8NEZ4, Q86X55, Q53H47, Q09028, Q6ZN18, Q15022, Q92800, Q16576, P26358, O43463, O14744, O14686, O00167, Q6PL18, Q53GL7, P06400, Q9NR48, Q92831, Q09472, O95696, Q8IXJ6 |
| organic hydroxy compound metabolic process | PPT1 | 22 | P42330, Q06520, P50225, P49888, P23141, P0DMM9, O43704, Q9UBM7, P14550, P13569, Q9NYA1, Q9NRA0, P31213, P18405, P19099, P15538, P05107, P14324, Q9NYB5, P17405, P11413, P00374 |
| negative regulation of transcription by RNA polymerase II | PPT2 | 22 | P84022, P10828, P24864, P24385, Q96T88, P40763, Q9Y6K1, Q9H5I1, Q6ZN18, Q15022, O75530, Q92800, P26358, O43463, Q92793, P09874, O75604, P06400, Q09472, P21675, Q9Y5X4, Q8IXJ6 |
| cell-substrate adhesion | PPT1 | 11 | P18084, P08514, P18564, P05556, P08648, P07237, P20701, P05107, P17301, P50281, P39900 |
| ion transmembrane transport | PPT1 | 35 | P05556, P03905, O43497, P13569, Q9NRA0, O15440, Q7RTT9, Q9Y694, Q96NT5, Q8TCC7, Q4U2R8, P41440, P15328, P14207, P11413, Q96SW2, P46098, P43681, O75762, P47869, Q16445, P18505, P18507, P47870, O00591, P34903, Q8N1C3, P78334, Q99928, P31644, O14764, P48169, P14867, P28472, Q9UN88 |
| macromolecule modification | PPT2 | 130 | P53350, Q8TD08, Q13627, P29317, Q9HCP0, Q8TDC3, Q5VT25, Q9P1W9, Q14012, O43293, Q7KZI7, Q13131, O14757, Q9BZL6, P31749, O14965, Q9UEE5, Q9H2X6, Q96SB4, Q8IW41, O75582, Q9HAZ1, Q13188, P51955, P51817, O96017, O96013, P27448, Q13153, P23443, Q9Y243, Q16513, P31751, Q96GD4, P17612, Q05655, P51946, O75116, P04637, Q8NEB9, P22612, Q8WWL7, O95067, P68400, O96020, P25440, P49768, P06748, P10415, Q8TEK3, O00141, P51812, O15530, P67870, P36507, P11274, P05771, P20248, Q02750, P14635, Q15078, P24864, P24385, Q13526, P78396, Q13563, Q8IV63, Q9H2K2, P78527, Q96T88, P40225, Q13490, Q15118, Q96B36, Q9Y6K1, Q9UMN6, Q9H5I1, Q99873, Q8WTS6, Q8NEZ4, Q86X55, Q53H47, Q09028, Q15022, O75530, Q92800, Q16576, P26358, O60678, O43463, O14744, O14686, P13500, P31947, O00571, O00167, Q92793, P18031, P09874, Q9H237, Q9Y6F1, Q9UGN5, Q9NR21, O95271, Q9H0J9, Q8N3A8, Q7Z3E1, Q53GL7, Q2NL67, P42574, O75604, P40189, P08887, P06400, Q9NR48, Q92831, Q09472, P21675, O95696, Q9HC16, O60725, Q53EL6, Q9NTG7, Q8IXJ6, P05067, Q6UB28, P50579, Q9UIQ6, P53582, P55786 |
| biological process involved in symbiotic interaction | PPT1 | 13 | P11142, P08238, P18084, P18564, P05556, P08648, P23284, P07237, P05362, P17301, P17405, O75351, P61073 |
| external encapsulating structure organization | PPT1 | 19 | P05556, P08253, P07477, Q9Y5R2, Q9UNA0, Q9NRE1, Q9NPA2, P51512, P51511, P50281, P45452, P39900, P22894, P09238, P09237, O75173, O60882, O14672, P27487 |
| signal transduction by p53 class mediator | PPT2 | 14 | O14757, P31749, O14965, Q9H2X6, Q8IW41, O96017, Q96GD4, P04637, P10415, O00141, O14744, O75604, Q09472, P21675 |
| synaptic transmission, GABAergic | PPT1 | 14 | P35348, P34972, P28222, P47869, Q16445, P18507, P47870, P34903, Q8N1C3, P78334, Q99928, P31644, P48169, P14867 |
| collagen catabolic process | PPT1 | 14 | P05556, P08253, Q9Y5R2, Q9NRE1, Q9NPA2, P51512, P51511, P50281, P45452, P39900, P22894, P09238, P09237, O60882 |
| collagen metabolic process | PPT1 | 15 | P05556, P08253, P17301, Q9Y5R2, Q9NRE1, Q9NPA2, P51512, P51511, P50281, P45452, P39900, P22894, P09238, P09237, O60882 |
| gamma-aminobutyric acid signaling pathway | PPT1 | 14 | P08908, P47869, Q16445, P18505, P18507, P47870, P34903, Q8N1C3, P78334, Q99928, P31644, P48169, P14867, P28472 |

**Table 3. KEGG pathways**

| **Description** | **protein_group** | **weight** | **protein_list** |
| --- | --- | --- | --- |
| Metabolic pathways | PPT2 | 47 | O00764, Q9Y2D0, Q9ULX7, Q8N1Q1, Q16790, P43166, P35218, P34913, P23280, P22748, P09917, P07451, P00915, P00403, O43570, O14684, P32320, P27707, P04183, O00142, P29474, Q9Y6K1, Q9H5I1, Q8WTS6, Q8NEZ4, Q53H47, Q92800, P26358, O43463, O14686, P23921, P36537, P19224, P48426, Q9NR48, Q9NTG7, Q8IXJ6, P30519, P28838, P15144, Q9UNK4, Q9NZK7, Q9NZ20, Q9BZM2, Q9BZM1, P14555, O15496 |
| Regulation of actin cytoskeleton | PPT1 | 12 | P18084, P08514, P18564, P05556, P08648, P20701, P05107, P17301, P61073, P62136, P36873, P61160 |
| GABAergic synapse | PPT1 | 17 | P46459, P47869, Q16445, P18505, P18507, P47870, O00591, P34903, Q8N1C3, P78334, Q99928, P31644, O14764, P48169, P14867, P28472, Q9UN88 |
| Taste transduction | PPT1 | 11 | P46098, P28566, P28222, P28221, P08908, P47869, Q16445, P34903, P31644, P48169, P14867 |
| Antifolate resistance | PPT1 | 11 | O15438, O15440, P31939, Q96NT5, Q05932, P41440, P22102, P15328, P14207, P04818, P00374 |
| Retrograde endocannabinoid signaling | PPT1 | 20 | P03905, O00519, Q9Y4D2, Q99685, P47869, Q16445, P18505, P18507, P47870, O00591, P34903, Q8N1C3, P78334, Q99928, P31644, O14764, P48169, P14867, P28472, Q9UN88 |
| Cell cycle | PPT2 | 17 | O96017, P51946, P04637, Q8WWL7, O95067, O96020, P20248, P84022, P14635, P24864, P24385, P78396, P78527, P31947, Q92793, P06400, Q09472 |
| Viral carcinogenesis | PPT2 | 18 | P17612, P04637, P22612, O96020, P20248, P62993, P24864, P24385, P78396, P40763, O00571, Q92793, P42574, Q9Y6K9, P40189, P06400, Q92831, Q09472 |
| HIF-1 signaling pathway | PPT2 | 14 | P23443, Q9Y243, P31751, P10415, P36507, P05771, Q02750, P40763, Q15118, P29474, Q92793, P08887, Q09472, P06730 |
| FoxO signaling pathway | PPT2 | 19 | Q13131, P31749, Q9Y243, P31751, Q8WWL7, O95067, O00141, O15530, P36507, Q02750, P84022, P14635, P62993, P24385, P40763, Q99873, Q8WTS6, Q92793, Q09472 |
| MicroRNAs in cancer | PPT2 | 19 | O75582, O96013, P04637, O96020, P10415, P36507, P05771, Q02750, P62993, P24864, P24385, O94925, P40763, Q9Y6K1, P26358, Q92793, P42574, Q09472, Q53EL6 |
| Neuroactive ligand-receptor interaction | PPT1 | 34 | P07477, Q9H228, Q99500, P21453, O95977, O95136, P43681, P35368, P35348, P34972, P34969, P28566, P28222, P28221, P21918, P08908, P47869, Q16445, P18505, P18507, P47870, O00591, P34903, Q8N1C3, P78334, Q99928, P31644, O14764, P48169, P14867, P28472, Q9UN88, P35030, P07478 |
| Apoptosis | PPT2 | 14 | Q9Y243, P31751, P04637, Q07817, P10415, O15530, P36507, Q02750, Q13490, P09874, Q9Y6F1, Q9UGN5, P42574, Q9Y6K9 |
| Kaposi sarcoma-associated herpesvirus infection | PPT2 | 14 | Q9Y243, P31751, P04637, Q8NEB9, P36507, Q02750, P24385, P40763, Q92793, P42574, Q9Y6K9, P40189, P06400, Q09472 |
| Human immunodeficiency virus 1 infection | PPT2 | 18 | O14757, P31749, O96013, Q13153, P23443, Q9Y243, P31751, Q8WWL7, O95067, Q07817, P10415, P36507, P05771, Q02750, P14635, P42574, Q9Y6K9, Q9HC16 |
| Human cytomegalovirus infection | PPT2 | 19 | P23443, Q9Y243, P31751, P17612, O75116, P04637, P22612, P36507, P05771, Q02750, P62993, P24385, P40763, P13500, P63092, P42574, Q9Y6K9, P08887, P06400 |
| Cellular senescence | PPT2 | 20 | O14757, P31749, Q9H2X6, O96017, Q9Y243, P31751, P04637, Q8WWL7, O95067, O96020, P36507, P20248, Q02750, P84022, P14635, P24864, P24385, P78396, Q09028, P06400 |
| p53 signaling pathway | PPT2 | 11 | O96017, P04637, O95067, O96020, Q07817, P10415, P14635, P24864, P24385, P31947, P42574 |
| Pancreatic cancer | PPT2 | 11 | P23443, Q9Y243, P31751, P04637, Q07817, Q02750, P84022, P24385, P40763, Q9Y6K9, P06400 |
| Colorectal cancer | PPT2 | 11 | P23443, Q9Y243, P31751, P04637, P10415, P36507, Q02750, P84022, P62993, P24385, P42574 |
| Hepatocellular carcinoma | PPT2 | 12 | P23443, Q9Y243, P31751, P04637, Q07817, P36507, P05771, Q02750, P84022, P62993, P24385, P06400 |
| Chronic myeloid leukemia | PPT2 | 12 | Q9Y243, P31751, P04637, Q07817, P36507, P11274, Q02750, P84022, P62993, P24385, Q9Y6K9, P06400 |
| EGFR tyrosine kinase inhibitor resistance | PPT2 | 12 | P23443, Q9Y243, P31751, Q07817, P10415, P36507, P05771, Q02750, P62993, P40763, P08887, P06730 |
| Wnt signaling pathway | PPT2 | 13 | P17612, O75116, P04637, P22612, P68400, P49768, P67870, P05771, P84022, P24385, Q92793, Q9H237, Q09472 |
| Neurotrophin signaling pathway | PPT2 | 13 | O75582, Q9Y243, P31751, Q05655, P04637, P49810, P49768, P10415, P51812, O15530, P36507, Q02750, P62993 |
| Growth hormone synthesis, secretion and action | PPT2 | 13 | Q9Y243, P31751, P17612, P22612, Q13936, P36507, P05771, Q02750, P62993, P40763, P63092, Q92793, Q09472 |
| JAK-STAT signaling pathway | PPT2 | 13 | Q9Y243, P31751, Q07817, P10415, P62993, P24385, P40225, P40763, Q92793, P40189, P42702, P08887, Q09472 |
| mTOR signaling pathway | PPT2 | 14 | Q13131, P31749, P23443, Q9Y243, P31751, O00141, P51812, O15530, P36507, P05771, Q02750, P62993, Q96B36, P06730 |
| Gastric cancer | PPT1 | 3 | P28702, P10826, Q15465 |
| Gastric cancer | PPT2 | 14 | P31749, P23443, Q9Y243, P31751, P04637, O96020, P10415, P36507, Q02750, P84022, P62993, P24864, P24385, P06400 |
| Nicotine addiction | PPT1 | 17 | P43681, P47869, Q16445, P18505, P18507, P47870, O00591, P34903, Q8N1C3, P78334, Q99928, P31644, O14764, P48169, P14867, P28472, Q9UN88 |
| Hepatitis B | PPT2 | 20 | Q9Y243, P31751, P04637, O96020, P10415, P36507, P05771, P20248, Q02750, P84022, P62993, P24864, P78396, P40763, O00571, Q92793, P42574, Q9Y6K9, P06400, Q09472 |
| Serotonergic synapse | PPT1 | 10 | P46098, P34969, P28566, P28222, P28221, P08908, P18505, P47870, P28472, P51589 |
| Serotonergic synapse | PPT2 | 12 | P17612, P22612, P18054, Q13936, O15296, P05771, Q02750, P09917, P63092, P42574, Q00975, P05067 |
| Morphine addiction | PPT1 | 19 | Q16445, P18505, P18507, P47870, O00591, P34903, Q8N1C3, P78334, Q99928, P31644, O14764, P48169, P14867, P28472, Q9UN88, Q08499, P27815, Q07343, Q08493 |
| Morphine addiction | PPT2 | 5 | P17612, P22612, P05771, P63092, Q00975 |

**Table 4. Compound list**

| **Drug** | **Cluster** | **Mechanism/Targets** | **Concentration (nM)** | **Solvent** | **Supplier Ref** | **Supplier** |
| --- | --- | --- | --- | --- | --- | --- |
| LY3009120 | 1 | pan-RAF inhibitor | 1-10000 | DMSO | HY-12558 | Medchem Express |
| Ruxolitinib | 1 | JAK1&2 inhibitor | 1-10000 | DMSO | CT-INCB-2 | ChemieTek |
| Birabresib | 1 | BET family inhibitor | 1-10000 | DMSO | S7360-2 | Selleck |
| Plicamycin | 1 | RNA synthesis inhibitor | 1-10000 | DMSO | sc-200909-7 | Santa Cruz Biotechnology |
| Teniposide | 2 | Topoisomerase II inhibitor | 1-10000 | DMSO | HY-13761 | Medchem Express |
| Silmitasertib | 2 | CSNK2A1 inhibitor | 1-10000 | DMSO | S2248 | Selleck |
| Ulixertinib | 2 | ERK inhibitor | 1-10000 | DMSO | CT-VRT752 | ChemieTek |
| Sapanisertib | 2 | mTOR1/2 Inhibitor | 0.1-1000 | DMSO | CT-INK128 | ChemieTek |

**Table 5. Patient characteristics**

| **Sample_ID** | **Diagnosis** | **Disease Stage** | **FAB** | **genetic Characteristics** | **age** | **Malignant Cell Percentage** |
| --- | --- | --- | --- | --- | --- | --- |
| AML_1 | C92.0 Acute myeloid leukaemia [AML] | Refractory | M5 | ["+8","+11"] | 67 | 59 |
| AML_2 | C92.0 Acute myeloid leukaemia [AML] | Relapse |  | ["Chromosomal abnormalities not checked"] | 75 | 49 |
| AML_3 | C92.0 Acute myeloid leukaemia [AML] | Diagnosis | M5 | ["No chromosomal abnormalities detected"] | 34 | 85 |
| AML_4 | C92.0 Acute myeloid leukaemia [AML] | Diagnosis | M2 | ["No chromosomal abnormalities detected"] | 44 | 55 |
| AML_5 | C92.0 Acute myeloid leukaemia [AML] | Relapse | M5 | ["No chromosomal abnormalities detected"] |  | 75 |
| AML_6 | C92.0 Acute myeloid leukaemia [AML] | Relapse | M0,FAB M1 | Not available |  | 83 |
| AML_7 | C92.0 Acute myeloid leukaemia [AML] | Relapse |  | ["inv(11)","add(14q)","inc[cp9]/46"] | 55 | 90 |
| AML_8 | C92.0 Acute myeloid leukaemia [AML] | Diagnosis | M2 | ["No chromosomal abnormalities detected"] | 75 | 90 |
| AML_9 | C92.0 Acute myeloid leukaemia [AML] | Diagnosis |  | ["No chromosomal abnormalities detected"] | 72 | 77 |
| AML_10 | C92.4 Acute promyelocytic leukaemia [PML] | Diagnosis | M3 | ["t(15;17)(q22;q12); PML-RARA*"] | 56 | 80 |
| AML_11 | C92.4 Acute promyelocytic leukaemia [PML] | Diagnosis | M3 | ["t(15;17)(q22;q12); PML-RARA*"] | 60 | 60 |
| AML_12 | C92.0 Acute myeloid leukaemia [AML] | Relapse |  | ["t(xx;11)(xx;q23) MLL-fusions"] | 7 | 100 |
| AML_13 | C92.0 Acute myeloid leukaemia [AML] | Diagnosis |  | ["t(8;21)(q22;q22); RUNX1-RUNX1T1","abn(11)"] | 72 | 86 |
| AML_14 | C92.0 Acute myeloid leukaemia [AML] | Diagnosis |  | ["No chromosomal abnormalities detected"] | 58 | 82 |
| AML_15 | C92.0 Acute myeloid leukaemia [AML] | Diagnosis |  | Not available | 64 | 53 |
| AML_16 | C92.0 Acute myeloid leukaemia [AML] | Diagnosis |  | Not available | 52 | 81 |
| Healthy_1 | Healthy |  |  |  |  |  |
| Healthy_2 | Healthy |  |  |  |  |  |
| Healthy_3 | Healthy |  |  |  |  |  |
| Healthy_4 | Healthy |  |  |  |  |  |
| Healthy_5 | Healthy |  |  |  | 66 |  |

**Table 6. Antibodies used in flow cytometry**

| **Biomarker** | **Fluorophore** | **Clone** | **iQue3 channel** | **Dilution** | **Manufacturer** | **Ref. No** |
| --- | --- | --- | --- | --- | --- | --- |
| CD38 | BV421 | HIT2 | VL1 | 1:800 | BD Biosciences | 562444 |
| CD11b | BV605 | ICRF44 | VL4 | 1:267 | BD Biosciences | 562721 |
| CD15 | BV786 | W6D3 | VL6 | 1:800 | BD Biosciences | 741013 |
| CD34 | PE | 563 | BL2 | 1:160 | BD Biosciences | 550761 |
| CD45 | V500 | HI30 | VL2 | 1:267 | BD Biosciences | 560777 |
| CD14 | APC | M5E2 | RL1 | 1:62 | BD Biosciences | 555399 |
| CD117 | PE/Cy7 | 104D2 | BL5 | 1:800 | Biolegend | 313212 |
| DRAQ7 | N/A | N/A | RL2 | 1:800 | BD Biosciences | 564904 |
| Annexin V | FITC | N/A | BL1 | 1:200 | BD Biosciences | 556419 |

**Table 7. FC results in different populations**

| **Drug Combinations** | **Sample** | **Live Ratio** | **Blast Ratio** | **Response** | **CD117**  **Ratio** | **Response** | **CD34**  **Ratio** | **Response** | **CD14**  **Ratio** | **Response** | **CD15**  **Ratio** | **Response** | **Imphocyte Ratio** | **Response** | **CD34+CD38-**  **Ratio** | **Response** | **CD34+CD38+**  **Ratio** | **Response** | **CD11b**  **Ratio** | **Response** |
| --- | --- | --- | --- | --- | --- | --- | --- | --- | --- | --- | --- | --- | --- | --- | --- | --- | --- | --- | --- | --- |

**LY3009120 &**

**Sapanisertib**

**LY3009120 &**

**Silmitasertib**

**LY3009120 &**

**Ulixertinib**

**Ruxolitinib & Silmitasertib**

**Ruxolitinib & Ulixertinib**

AML_3

Average

0.84

0.65

0.76

0.86

32.57

47.01

0.48

0.25

33.22

36.57

0.59

0.23

33.84

48.61

0.25

0.09

21.19

43.91

0.32

0.20

28.75

52.85

0.03

0.03

3.45

21.37

0.55

0.18

22.96

21.52

0.25

0.12

52.35

58.55

0.18

0.07

48.06

56.54

**Total average**

0.65

0.69

35.70

0.26

30.96

0.30

34.88

0.17

56.47

0.24

54.75

0.06

13.75

0.25

35.11

0.09

55.24

0.11

55.60

AML_4	0.83	0.94	42.7	0.24	0	0.06	30.97	0	33.19	0.11	47.08	0.03	1.75	0.00	0.00	0.05	41.44	0.02	38.84

AML_1	0.77	0.16	53.37	0.05	39.7	0.03	42.75	0.54	74.98	0.35	70.83	0.04	5.65	0.03	69.47	0.07	80.53	0.29	77.14

AML_2	0.56	0.76	23.53	0.49	14.44	0.75	26.7	0	89.93	0.23	77.18	0.19	9.44	0.67	25.99	0.04	70.77	0.02	62.87

Average	0.72	0.56	39.49	0.34	29.47	0.46	37.61	0.26	79.50	0.30	71.78	0.09	7.77	0.42	46.72	0.12	70.65	0.16	71.34

AML_3	0.84	0.76	41.56	0.48	34.26	0.59	43.38	0.25	73.6	0.32	67.32	0.03	8.21	0.55	44.70	0.25	60.64	0.18	74.00

AML_2	0.56	0.76	13.48	0.49	10.01	0.75	14.08	0	81.54	0.23	61.66	0.19	4.94	0.67	13.05	0.04	45.12	0.02	41.56

AML_4	0.83	0.94	27.09	0.24	27.71	0.06	5.1	0	28.26	0.11	22.23	0.03	3.45	0.00	20.70	0.05	6.99	0.02	25.11

AML_1	0.77	0.16	21.41	0.05	18.12	0.03	17.2	0.54	45.28	0.35	41.73	0.04	5.42	0.03	51.78	0.07	52.72	0.29	48.89

Average	0.72	0.62	20.66	0.26	18.61	0.28	12.13	0.18	51.69	0.23	41.87	0.09	4.60	0.24	28.51	0.05	34.94	0.11	38.52

AML_2	0.56	0.76	11.64	0.49	6.33	0.75	12.93	0	77.35	0.23	64.3	0.19	3	0.67	12.10	0.04	52.39	0.02	45.32

AML_1	0.77	0.16	24.86	0.05	33.82	0.03	31.8	0.54	52.97	0.35	48.24	0.04	2.79	0.03	53.36	0.07	56.09	0.29	52.93

AML_5	0.27	0.89	46.47	0.02	68.86	0.04	51.27	0.03	52.94	0.16	55.41	0.02	44.94	0.00	61.06	0.04	50.26	0.02	56.25

Average	0.53	0.60	27.66	0.19	36.34	0.27	32.00	0.19	61.09	0.25	55.98	0.08	16.91	0.23	42.17	0.05	52.91	0.11	51.50

AML_5	0.27	0.89	66.46	0.02	75.83	0.04	80.45	0.03	74.44	0.16	82.02	0.02	53.01	0.00	78.37	0.04	80.50	0.02	77.02

AML_4	0.83	0.94	41.86	0.24	0.85	0.06	32.66	0	31.43	0.11	38.44	0.03	2.26	0.00	2.52	0.05	41.21	0.02	44.50

AML_3	0.84	0.76	31.25	0.48	29.04	0.59	29.29	0.25	23.17	0.32	31.89	0.03	4.76	0.55	18.88	0.25	58.15	0.18	59.55

Average	0.65	0.86	46.52	0.25	35.24	0.23	47.47	0.09	43.01	0.20	50.78	0.03	20.01	0.18	33.25	0.12	59.95	0.07	60.36

AML_5	0.27	0.89	65.75	0.02	76.48	0.04	81.02	0.03	77.35	0.16	82.71	0.02	58.9	0.00	41.59	0.04	81.87	0.02	82.72
